# Can genome-based analysis be the new gold-standard for routine Salmonella serotyping?

**DOI:** 10.1101/806208

**Authors:** Sangeeta Banerji, Sandra Simon, Andreas Tille, Antje Flieger

## Abstract

*Salmonella enterica* is the second most reported bacterial cause of food-borne infections in Europe. Therefore molecular surveillance activities based on pathogen subtyping are an important measure of controlling *Salmonellosis* by public health agencies. In Germany, at the federal level, this work is carried out by the National Reference Center for *Salmonella* and other Bacterial Enteric Pathogens (NRC). With rise of next generation sequencing techniques, the NRC has introduced whole-genome-based typing methods for *S. enterica* in 2016. In this study we report on the feasibility of genome-based *in silico* serotyping in the German setting using raw sequence reads. We found that SeqSero and seven gene MLST showed 98% and 95% concordance, respectively, with classical serotyping for the here evaluated serotypes, including the most common German serotypes *S.* Enteritidis and *S.* Typhimurium as well as less frequently found serotypes. The level of concordance increased to >99% when the results of both *in silico* methods were combined. However, both tools exhibited misidentification of monophasic variants, in particular monophasic *S.* Typhimurium and therefore need to be fine-tuned for reliable detection of this epidemiologically important variant. We conclude that with adjustments *Salmonella* genome-based serotyping might become the new gold standard.

## Introduction

Subtyping of bacterial enteric pathogens, such as *Salmonella enterica*, traditionally relies on serotyping. The species *Salmonella enterica* is divided into six subspecies and consists of more than 2600 serovars, which are classified according to the White-Kauffmann-Le Minor Scheme ^1^. Serotyping is based on determination of somatic O antigens and flagellin H antigens by reaction with specific antisera. Most *S. enterica* serovars have two alternately expressed H antigens, also referred to as ‘phases’. The phase-1 and phase-2 flagellin proteins are encoded by *fliC* and *fljB*, respectively. The phase switch is regulated by the invertase *hin* and the *fliC* repressor gene *fljA* ^*2*^. Therefore, the specific antigenic formula consists of three positions: the first position represents the O antigens, the second and third positions the two different flagellin H antigens. Each antigen position is separated by a colon, i.e. O:H1:H2. The antigenic formula for *S.* Typhimurium for example is accordingly 1,4,[5],12:i:1,2. There are variants of *S.* Typhimurium, which express only one flagellin and which therefore are referred to as monophasic *S*. Typhimurium. *S.* Enteritidis on the other hand does not possess a second flagellin per se, which is reflected in the antigenic formula: 1,9,12:g,m:-. It should be noted that some serovars share the same antigenic formula and require additional testing for unambiguous identification, e.g. the clinically important serovar *S.* Chloeraesuis shares its antigenic formula 6,7:c:1,5 with serovars *S.* Paratyphi C and *S.* Typhisuis. A differentiation is possible based on biochemical characteristics or PCR ^3^.

With rise of next generation sequencing (NGS) techniques, genomic typing tools have become increasingly popular and effective. Several *in silico* classification tools employing NGS data are available for *Salmonella*. The serotyping tools are either based on identifying and characterizing the serotype-determining genes or derive the serotype from *in silico* Multi Locus Sequence Typing (MLST) or a combination of both methods. MLST-based serotyping was sparked by the observation of Achtmann et al. that the phylogeny derived from MLST sequence types correlates with serotypes ^4,5^. Achtmann and his group are also the developers of Enterobase, a platform for the phylogenetic analysis of selected bacteria, including *Salmonella* ^6^. A report of the Establishing Next Generation Sequencing Ability for Genomic Analysis in Europe (ENGAGE) consortium identified four serotyping tools, specifically Metric-Oriented Sequence Typer (MOST), SeqSero, *Salmonella*TypeFinder and SISTR, which were benchmarked for their performance and were found to have correlation rates between 65% and 88% with classical serotyping (http://www.engage-europe.eu/resources/benchmarking). MOST is a pipeline developed and employed by Public Health England, which infers an MLST type with a modified version of the program SRST, which was developed for deducing a sequence type from short reads, and utilizes a local database for identification of corresponding serotypes ^7,8^. SeqSero is an *in silico* serotyping program and determines the presence of O and H antigen loci within the NGS data, which correspond to the antigens involved in classical serotyping ^9^. *Salmonella*TypeFinder is a pipeline developed by the Danish Technical University, which runs SeqSero and determines the MLST type using an in-house MLST calling tool, and then both results are used for determination of the serotype (https://bitbucket.org/genomicepidemiology/salmonellatypefinder/src/master/). Another typing platform is SISTR, which predicts the serotype by a combination of *in silico* hybridization and extended MLST, incorporated into a ‘Microbial *in Silico* Typing’ engine ^10^.

Classification of *Salmonella* by serotyping is especially important for epidemiological investigations and is often routinely performed in its full scheme at National Reference Centers or Laboratories. It is also implemented at the German National Reference Center for *Salmonella* and other Bacterial Enteric Pathogens (NRC). The NRC receives around 3,000-4,000 *Salmonella* isolates per year from human infections for further characterization. The most common serotypes submitted are *S.* Enteritidis and *S.* Typhimurium, followed by other broad host range serotypes like *S.* Infantis and *S.* Derby ^11^. Since 2016 the NRC has been shifting towards NGS-based analysis ^12^.

Our aim for this study was to estimate whether NGS-based serotyping was feasible as a means of replacing traditional serotyping in our setting. The success rate of classical serotyping depends on many factors (e.g. access to high quality antisera, training of staff, and experience with rare serotypes) and was found to average worldwide at 82% and for European countries at 89% correct results in 2007 ^13^. Whereas the O antigens are determined within a few hours, characterization of the H phases may require up to 7 days. If the NRC replaced classical serotyping with a genome-based *in silico* typing method, this method should ideally match the high reported success rate of classical serotyping^13^. Genome-based typing tools have performed well in several studies with maximum reported concordance levels of approximately 92% for SeqSero ^9^ and approximately 94 % for SISTR 10,14. However, previous studies used assembled genomes for *in silico* typing. Only very recently, Ibrahim and Morin also reported results obtained with paired reads using the web-based application of SeqSero 1.0 ^15^. Genome assembly requires additional time and computing resources, which is a drawback for routine analysis of a large number of genomes.

Our goal for this study was therefore to directly use raw reads in order to save time and computing resources. Thus, our requirements for the tools were that the input data should need minimal preprocessing and should potentially fit into our existing analysis pipeline (Ridom SeqSphere^+^) ^16^. Since we need to process a large number of sequences, offline availability was also of major importance. SeqSero fulfilled all of these requirements (when used as a command line tool). The other above mentioned tools did not as they either use different allele detection algorithms for determination of MLST sequence types than Enterobase (MOST and *Salmonella*TypeFinder) or require an assembled genome (SISTR). Therefore we decided to assess the performance of SeqSero and the Enterobase MLST scheme from Achtman et al. for serotype prediction ^4^.

## Results

The aim of this study was to assess two *in silico* serotype prediction tools, namely SeqSero and MLST via SeqSphere/Enterobase for their performance in routine *Salmonella* typing at the NRC. We chose 520 *Salmonella* isolates, mainly of human origin and predominantly from the years 2014-2018 as the data set for analysis. The selection comprised very frequently found serotypes as well as less frequent serotypes (Table 1). We investigated a total of 20 different serotypes and also looked at monophasic variants as well as rough phenotypes.

**Table 1.** Overview of serotype prediction with SeqSero and MLST. Serotype was first determined by classical serotyping. Whole genome sequences were then analyzed with SeqSero or MLST. Correlation means that the predicted serotype was the same as the classically determined serovar. Ambiguous means that the correct serotype was listed among others. Prediction failure means that no complete antigenic formula was derived. Miscorrelation means that a wrong antigenic formula was derived. Overall (%) is the same as Total (%) except that Ambiguous (%) is added to Correlation (%). Final results are shown, i.e. after resequencing if data quality was not met.

| Serotype | Sequen-<br>ced<br>Isolates | Correlation |  |  | Ambiguous |  |  | Prediction failure |  |  | Miscorrelation |  |  |
| --- | --- | --- | --- | --- | --- | --- | --- | --- | --- | --- | --- | --- | --- |
|  |  | Seq-Sero | MLST | Seq-Sero<br>+<br>MLST | Seq-Sero | MLST | Seq-Sero<br>+<br>MLST | Seq-Sero | MLST | Seq-Sero<br>+<br>MLST | Seq-Sero | MLST | Seq-Sero<br>+<br>MLST |
| Agona | 3 | 3 | 3 | 3 | 0 | 0 | 0 | 0 | 0 | 0 | 0 | 0 | 0 |
| Choleraesuis | 33 | 0 | 33 | 33 | 30 or<br>Typhisuis<br>or<br>Paratyphi C | 0 | 0 | 3 | 0 | 0 | 0 | 0 | 0 |
| Choleraesuis<br>monophasic | 3 | 0 | 0 | 0 | 0 | 0 | 0 | 0 | 0 | 0 | 3 | 3 | 3 |
| Derby | 55 | 55 | 55 | 55 | 0 | 0 | 0 | 0 | 0 | 0 | 0 | 0 | 0 |
| 11:z41:e,n,z15<br>(novel serovar) | 10 | 10 | 10 | 10 | 0 | 0 | 0 | 0 | 0 | 0 | 0 | 0 | 0 |
| Enteritidis | 115 | 115 | 115 | 115 | 0 | 0 | 0 | 0 | 0 | 0 | 0 | 0 | 0 |
| Infantis | 50 | 49 | 50 | 50 | 0 | 0 | 0 | 1 | 0 | 0 | 0 | 0 | 0 |
| Kentucky | 7 | 7 | 7 | 7 | 0 | 0 | 0 | 0 | 0 | 0 | 0 | 0 | 0 |
| Kintambo | 3 | 0 | 3 | 3 | 3 or<br>Washington | 0 | 0 | 0 | 0 | 0 | 0 | 0 | 0 |
| Kottbus | 12 | 0 | 12 | 12 | 12 or<br>Ferruch | 0 | 0 | 0 | 0 | 0 | 0 | 0 | 0 |
| Mbandaka | 15 | 15 | 15 | 15 | 0 | 0 | 0 | 0 | 0 | 0 | 0 | 0 | 0 |
| Mikawasima | 10 | 10 | 10 | 10 | 0 | 0 | 0 | 0 | 0 | 0 | 0 | 0 | 0 |
| Muenchen | 25 | 0 | 25 | 25 | 25 or<br>Virginia | 0 | 0 | 0 | 0 | 0 | 0 | 0 | 0 |
| Paratyphi B | 6 | 6 | 6 | 6 | 0 | 0 | 0 | 0 | 0 | 0 | 0 | 0 | 0 |
| Paratyphi B<br>monophasic | 1 | 1 | 1 | 1 | 0 | 0 | 0 | 0 | 0 | 0 | 0 | 0 | 0 |
| Paratyphi C | 2 | 0 | 2 | 2 | 2 or<br>Cholerae-<br>suis or<br>Typhisuis | 0 | 0 | 0 | 0 | 0 | 0 | 0 | 0 |
| Poano | 2 | 2 | 2 | 2 | 0 | 0 | 0 | 0 | 0 | 0 | 0 | 0 | 0 |
| Strathcona | 2 | 2 | 2 | 2 | 0 | 0 | 0 | 0 | 0 | 0 | 0 | 0 | 0 |
| Stourbridge | 14 | 14 | 14 | 14 | 0 | 0 | 0 | 0 | 0 | 0 | 0 | 0 | 0 |
| Sundsvall | 1 | 0 | 1 | 1 | 1 or<br>Soahanina<br>or Sundvall | 0 | 0 | 0 | 0 | 0 | 0 | 0 | 0 |
| Typhi | 74 | 74 | 74 | 74 | 0 | 0 | 0 | 0 | 0 | 0 | 0 | 0 | 0 |
| Typhimurium<br>biphasic | 52 | 52 | 32 | 52 | 0 | 0 | 0 | 0 | 0 | 0 | 0 | 20 | 0 |
| Typhimurium<br>monophasic | 19 | 17 | 17 | 19 | 0 | 0 | 0 | 0 | 0 | 0 | 2 | 2 | 0 |
| Serologically<br>rough | 6 | 5 | 6 | 6 | 0 | 0 | 0 | 1 | 0 | 0 | 0 | 0 | 0 |
| Total number | 520 | 437 | 495 | 517 | 73 | 0 | 0 | 5 | 0 | 0 | 5 | 25 | 3 |
| Total (%) | 100.0 | 84.0 | 95.2 | 99.4 | 14.0 | 0 | 0 | 1.0 | 0 | 0 | 1.0 | 4.8 | 0.6 |
| Overall (%) | 100.0 | 98.0 | 95.2 | 99.4 | - | - | - | 1.0 | 0 | 0 | 1.0 | 4.8 | 0.6 |

### Data quality is an important bottleneck

Initially, we did not set a quality threshold for the raw read sequence files. In the course of the analysis we noticed that analysis with SeqSero 1.0 and/or Ridom SeqSphere^+^ failed if the file sizes of the raw sequence reads were lower than average (<50,000 KB). Since Zhang et al. only included data for analysis with SeqSero 1.0 with a minimal coverage of 10-fold, we aimed for the same quality threshold ^9^. Given an average genome size of approximately 4.8 Mb *for Salmonella enterica subsp. enterica*, we calculated that a theoretical coverage of ≥10-fold could only be achieved by a minimal read number of 100,000 each for paired end reads with a theoretical read length of 250: theoretical coverage = total number of reads x length of each read [bp] / genome size [bp]. Sequencing was repeated for cases not meeting the minimal read number (Fig. S1).

### SeqSero analysis correctly predicted the serovar in 98% of the isolates

SeqSero 1.0 predicted the serotype in 84% of analyzed strains in accordance to the classical serovar. In additional 14 % of the cases the antigenic formula was shared by more than one serovar and SeqSero 1.0 predicted all eligible serotypes, e.g. Choleraesuis, Typhisuis or Paratyphi C for the antigenic formula 6,7:c:1,5, which we rated as ambiguous. These cases require additional testing as they would if determined by classical serotyping. Therefore an ambiguous prediction was counted as a correlating result in the overall summary. The total rate of correlation (correlation + ambiguous prediction) with our laboratory results was therefore 98% (Table 1). Five cases of prediction failure (1%) occurred, all of which involved failed prediction of the O-7 antigen. Additionally, five cases (1%) of miscorrelation were found, which concerned monophasic strains of *S.* Typhimurium and *S.* Choleraesuis (Table 1).

### Monophasic variants are only predicted correctly if they lack the flagellin genes

17 out of 19 monophasic *S.* Typhimurium strains were correctly predicted by SeqSero 1.0 using raw reads (Table 1). In two cases, SeqSero 1.0 predicted phenotypically monophasic *S.* Typhimurium as biphasic. In order to investigate this discrepancy we analyzed the respective whole genome sequences by *de novo* assembly. The isolate ERR2003330 lacked approximately 250 nucleotides in the central part of the *fljB* gene as well as the whole *hin* gene (Fig. S2). Expression of the phase-2 flagellin gene *fljB* is co-regulated by the invertase gene *hin* and the *fliC* repressor gene *fljA* ^2^. Apparently a transposase, *tnpA*, had integrated into this region. This explains why the second phase could not be detected by classical serotyping. Since SeqSero 1.0 only checks whether the *fliC* and *fljB* alleles are present, it would explain why the lack of the *hin* gene was not detected by the program and the partial deletion of the *fljB* gene might have been too small to be detectable when using raw reads. We noted that SeqSero 1.0 correctly predicted the isolate to be monophasic when the analysis was performed with a5-assembled contigs. During preparation of this manuscript a new version of SeqSero called SeqSero 2.0 was available from github (https://github.com/denglab/SeqSero 2.0) and we rechecked the two non-correlating results with SeqSero 2.0 in the default k-mer-based mode. The program correctly classified isolate ERR2003330 as monophasic, probably due to the partial deletion in the *fljB* gene. However, when we used SeqSero 2.0 with a5 assembled contigs it classified the isolate wrongly as biphasic *S.* Typhimurium. The second isolate ERR2003327 had a transposon integrated into the *fljB* gene most probably rendering it non-functional. This isolate was identified to be biphasic with both SeqSero 1.0 and the k-mer-based approach of SeqSero 2.0 when using raw reads, because the *fljB* gene is fully present but interrupted. When using SeqSero 1.0 and SeqSero 2.0 with a5 assembled contigs isolate ERR2003327 was correctly predicted to be monophasic by both versions of the program.

Phenotypically monophasic variants of other serovars harboring *fliC* and *fljB* were also not recognized by SeqSero 1.0 or SeqSero 2.0, in particular three strains of *S.* Choleraesuis var. Kunzendorf not expressing phase-1 flagellum gene *fliC* (ERR3264001, ERR3264026, ERR3264035). This corroborates the fact that phenotypic traits are sometimes difficult to detect by *in silico* measures. A monophasic variant of *S.* Paratyphi B variant Java was recognized as monophasic *S.* Paratyphi B by SeqSero 1.0 and was additionally recognized to be L(+)-tartrate positive by SeqSero 2.0.

### Serovar prediction of rough strains is possible by means of SeqSero

Importantly, SeqSero 1.0 was able to predict a serotype for five out of six isolates with a rough phenotype, where classical serotyping was not successful (ERR3263893: *S.* Typhimurium, ERR3263889: *S.* Typhimurium monophasic, ERR3263894 [*S. e.* subspecies II]: 58:z6:z39, ERR3263880: *S.* Typhimurium monophasic, and ERR3263875: *S.* Typhimurium monophasic). We classified this as a correlation. For the rough strain ERR3264036 SeqSero 1.0 did not generate a full antigenic formula. PCR analysis according to Woods et al. 2008 targeting a 12.8-kb region specific to *S. Choleraesuis* yielded the serotype Choleraesuis ^3^. In this case, SeqSero 1.0 was only able to predict a partial antigenic formula for Choleraesuis (-:c:1,5). SeqSero 2.0 was likewise not able to provide a complete antigenic formula for this particular strain. When we mapped the raw reads against the respective *wzy* allele (locus tag EL48_RS10980), we found the allele and the surrounding region (EL48_RS10955-EL48_RS11010) missing (Fig. S3). We conclude that the rough phenotype of this particular isolate had a genetic basis.

### SeqSero does not reliably predict the O-7 antigen

We found five cases of prediction failure when using SeqSero 1.0 and all five cases involved failed prediction of the O-7 antigen, which is part of the epidemiologically important serovars *S.* Choleraesuis and *S.* Infantis (ERR3264036, ERR3264076, ERR3264063, ERR3264067, and ERR3264066). Except for Isolate ERR3264036, the remaining four cases had an intact *wzy* allele but only few reads mapped to the O-7 locus (Fig. S4 and S5). When we performed the analysis with SeqSero 2.0 the k-mer based approach yielded a complete antigenic formula which correlated with the laboratory phenotypes for all cases except for the rough strain ERR3264036, where the *wzy* allele is missing.

### SeqSero is well suited for routine high-throughput analysis of raw reads with the exception of atypical monophasic strains

In summary, SeqSero 1.0 is an easy to use tool, which is available as free software from the website of the developers, or as an official Debian package from the Debian website. Currently an alpha test version of SeqSero 2.0 is available on Github with additional features, e.g. k-mer based approach and integrated identification of the taxonomic ID with SalmID in the allele based mode for subspecies identification of ambiguous serovars. When using SeqSero 1.0 with Illumina paired end raw reads we achieved a correlation rate of 98%. The reasons for initial miscorrelation were mainly low data quality, which could be resolved by repeating the sequencing (Fig. S1). SeqSero 1.0 was able to predict a serotype for all rough isolates, except one. It correctly predicted monophasic variants if the flagellin genes *fliC* and/or *fljB* were missing. However, if the flagellin genes were only disrupted and/or other genes required for flagellar expression / phase transition were missing, SeqSero 1.0 and SeqSero 2.0 were not always able to reliably recognize monophasic variants. We conclude that with the exception of atypical monophasic variants of *S.* Typhimurium and other serovars and genetically rough strains (i.e. lack of O antigen determining genes) SeqSero is able to correctly predict the vast majority of common serovars circulating in Germany.

### MLST analysis correctly predicted the serovar in 95% of the isolates

MLST predicted the serotype in 95% of *Salmonella* isolates in concordance to the classical serovar found by serotyping (Table 1). Notably, all six rough isolates were assigned to a sequence type and a corresponding serotype. The prediction differed in 25 cases (5%) from the phenotypic classification all of which involved second phase miscorrelation. Figure 1 shows an UPGMA (unweighted pair group method with arithmetic mean) Tree based on MLST and color coded according to the serovar obtained by slide agglutination. As expected, there is a clear correlation between serotype and one or more closely related STs for the majority of isolates (Fig. 1). *S.* Enteritidis for example is distributed into the two closely related STs: ST 11 and ST 183. The *S.* Typhi isolates of our collection spread across five different but closely related STs: ST 1, ST2, ST 3677, ST 2173 and ST 2209 (Fig. 1).

**Fig.1.**
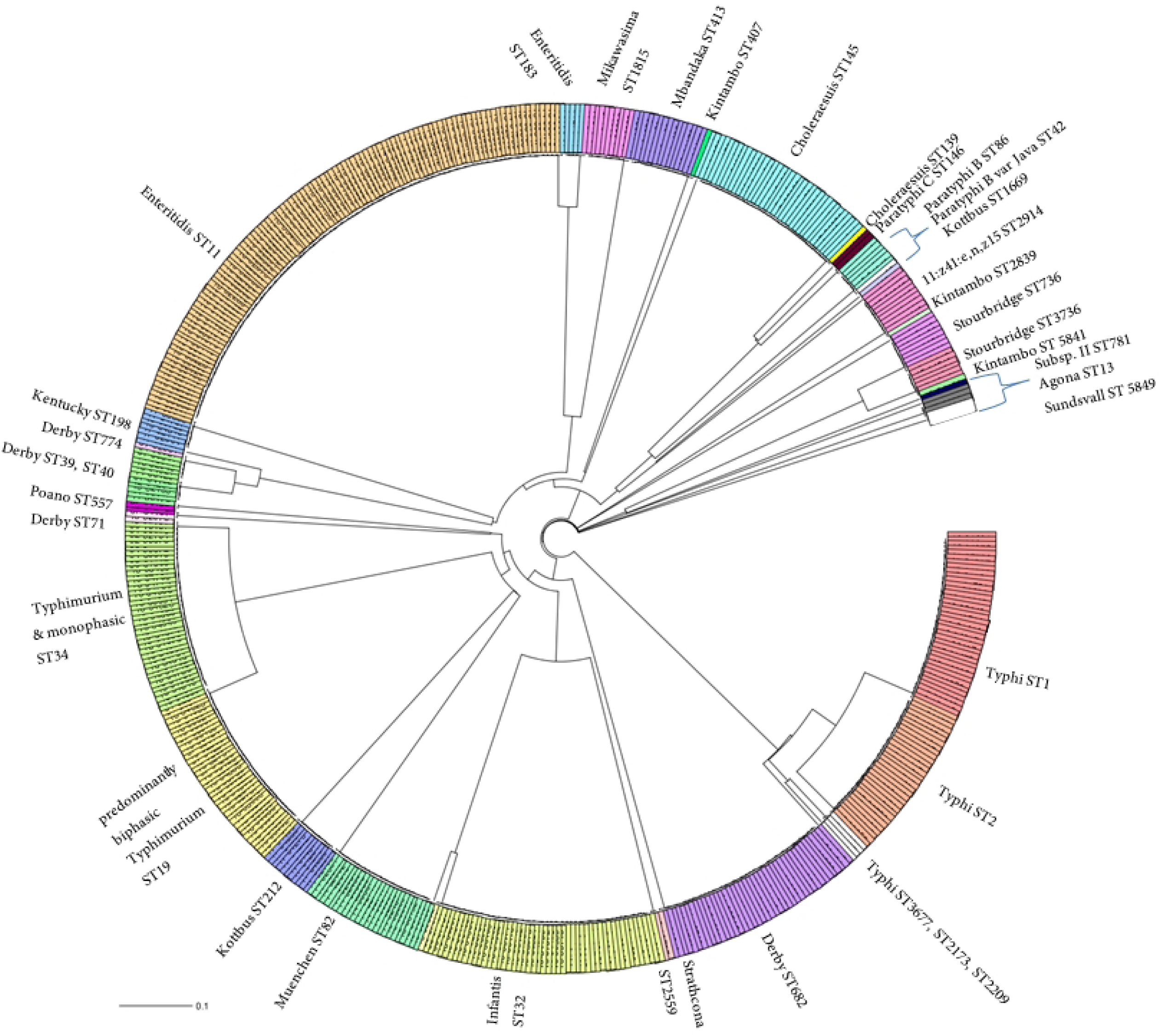
Unweighted Pair Group Method with Arithmetic mean (UPGMA) tree of all investigated isolates based on 7-gene MLST. The tree shows that serovars correlate to STs. Colors are based on ST.

### Sequence types do not consistently correlate with detection of flagellin antigens

It is notable that for the majority of isolates in Enterobase the antigenic formula is not provided by the user. Nevertheless, the majority of Enterobase strains belonging to ST 34, which had an antigenic formula provided, represented monophasic *S.* Typhimurium (203 out of 209 isolates as of May 2019). Therefore we assigned all ST 34 strains to monophasic Typhimurium. Enterobase strains belonging to ST 19 were a mix of monophasic and biphasic Typhimurium. We opted to classify all ST 19 isolates as biphasic Typhimurium although this would result in a high error rate. We preferred this to no classification at all. We obtained correlating results between MLST and classical serotyping for 17 out of 19 (89.5%) of our monophasic *S.* Typhimurium strains. Only 32 out of 52 biphasic *S.* Typhimurium belonged to ST 19 (61.5%) and were therefore also classified as biphasic with MLST. 20 out of 52 (38.5%) phenotypically biphasic Typhimurium belonged to ST 34 and were therefore wrongly classified as monophasic by MLST. We also checked whether the classification of monophasic and biphasic *S.* Typhimurium would be improved by clustering according to core genome MLST. Figure 2 depicts a minimum spanning tree of only monophasic and biphasic *S.* Typhimurium isolates (including three rough isolates) based on the Enterobase core genome MLST scheme. The isolates cluster according to their ST rather than to their flagellin expression.

**Fig.2.**
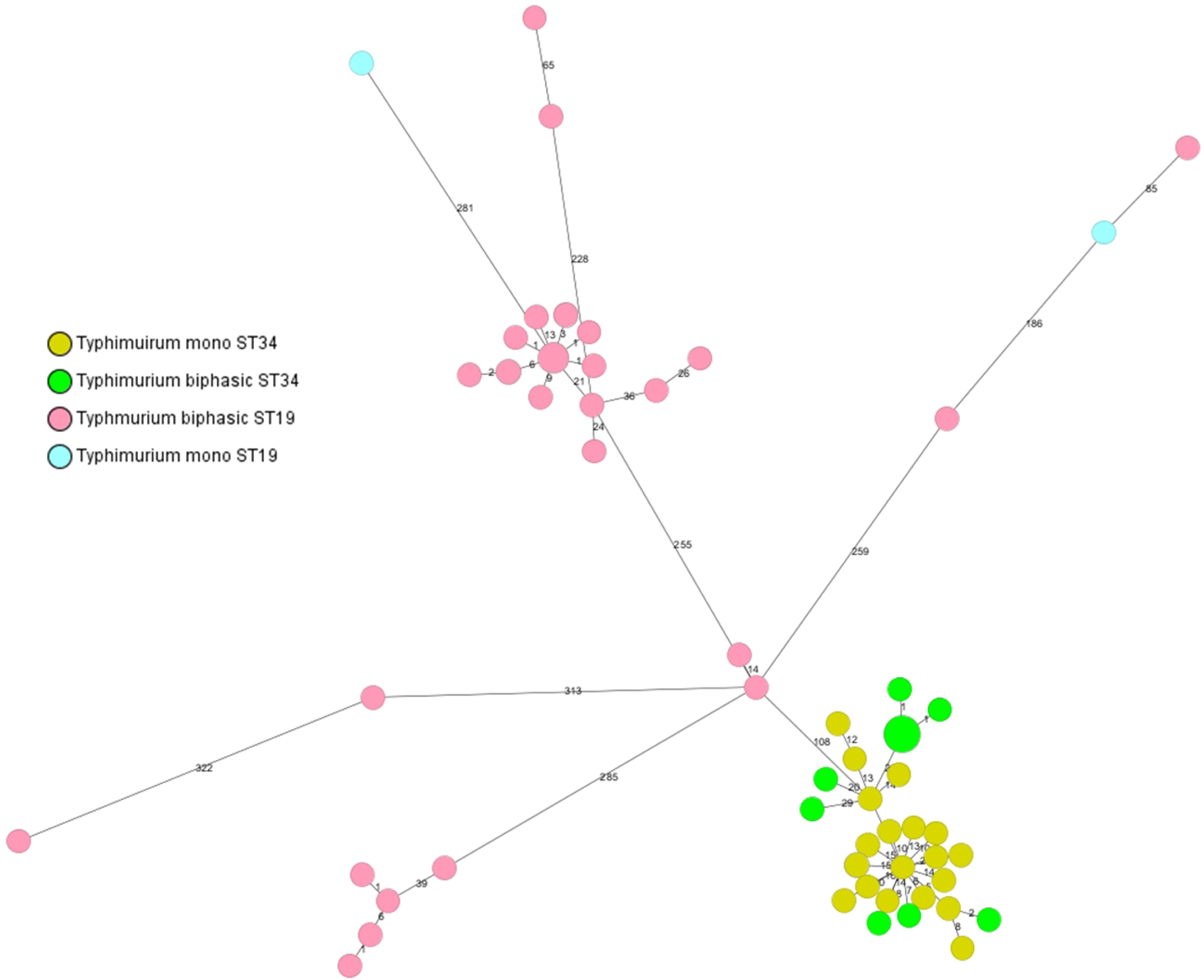
Minimal Spanning tree of monophasic and biphasic Typhimurium isolates based on the Enterobase core genome MLST scheme and 7-gene MLST. The tree reveals that *S.* Typhimurium isolates cluster according to ST rather than expression of flagellin. Colors are based on phase and STs.

The *S.* Choleraesuis isolates of our collection, phenotypically lacking FliC were also not correctly classified by MLST typing. In Enterobase monophasic *S.* Choleraesuis var. Kunzendorf predominantly belonged to ST 66, whereas our isolates belonged to ST 145. Interestingly, MLST distinguished the monophasic *S.* Paratyphi B var. Java as such, since ST 42 mostly consists of monophasic var. Java entries in Enterobase.

### MLST-based serotype prediction additionally provides phylogenetic context

The majority of our serotypes could each be assigned to a single eBG: e.g. *S.* Typhimurium to eBG 1, *S.* Enteritidis to eBG 4, *S.* Typhi to eBG 13 and *S.* Choleraesuis to eBG 6 (Table 2). This is also reflected in the phylogenetic tree, where the different STs, which comprise the same serovar and belong to the same eBG are located in the same branch (Fig. 1). This indicates that German strains belonging to these serovars stem from a common ancestor ^4,17^. One advantage of MLST serotype prediction compared to SeqSero was that there was no ambiguous serotype prediction. Different serovars with the same antigenic formula split into distinct eBurst groups (e.g. *S.* Choleraesuis eBG 6 and *S.* Paratyphi C eBG 20). MLST additionally provided important phylogenetic information, e.g. the *S.* Derby strains in our collection were of a polyphyletic nature as they split into three different eBGs (Table 2 and Fig. 1). In conclusion, MLST-based serotype prediction also proved to be very successful with the draw-back of not being able to distinguish between monophasic and biphasic *S.* Typhimurium as well as between *S.* Choleraesuis and monophasic *S.* Choleraesuis var. Kunzendorf.

**Table 2.** Overview of Serovars with corresponding MLST sequence types and e-Burst groups

| ***Salmonella* Serotype (Enterobase)** | **Sequence type** | **e-Burst Group** | **Number of Isolates** |
| --- | --- | --- | --- |
| Agona | 13 | 54 | 3 |
| Choleraesuis | 139 | 6 | 1 |
| Choleraesuis | 145 | 6 | 36 |
| Derby | 39 | 57 | 6 |
| Derby | 774 | 57 | 1 |
| Derby | 40 | 57 | 5 |
| Derby | 71 | 244 | 2 |
| Derby | 682 | 264 | 41 |
| Enteritidis | 11 | 4 | 110 |
| Enteritidis | 183 | 4 | 5 |
| 11:z41:e,n,z15 | 2914 | 472 | 10 |
| Infantis | 32 | 31 | 49 |
| Infantis | 2283 | 31 | 1 |
| Kentucky | 198 | 56 | 7 |
| Kintambo | 407 | 400 | 1 |
| Kintambo | 2839 | ND | 1 |
| Kintambo | 5841 | ND | 1 |
| Kottbus | 212 | 64 | 11 |
| Kottbus | 1669 | 63 | 1 |
| Mikawasima | 1815 | 247 | 10 |
| Mbandaka | 413 | 62 | 15 |
| Muenchen | 82 | 8 | 25 |
| Paratyphi B | 86 | 5 | 6 |
| Paratyphi B mono (var Java) | 42 | 32 | 1 |
| Paratyphi C | 146 | 20 | 2 |
| Poano | 557 | 87 | 2 |
| Strathcona | 2559 | ND | 2 |
| Stourbridge | 736 | 438 | 8 |
| Stourbridge (only RKI data) | 3736 | 464 | 6 |
| Sundsvall (first typed as Poano) | 488 | 305.2 | 1 |
| Subsp. II | 781 | 340 | 1 |
| Typhi | 1 | 13 | 38 |
| Typhi | 2 | 13 | 32 |
| Typhi | 2173 | 13 | 1 |
| Typhi | 2209 | 13 | 1 |
| Typhi | 3677 | 13 | 2 |
| Typhimurium & monophasic var. | 19 | 1 | 36 |
| Typhimurium & monophasic var. | 34 | 1 | 39 |
| Total | 39 | >26 | 520 |

### Combination of SeqSero and MLST increases robustness of prediction

After performing both analyses independently, we combined SeqSero 1.0 and MLST and used both results for predicting the serotype. In general, there was good agreement between the two methods. In case of disagreement, we evaluated the sequences individually. There were 24 cases of disagreement between SeqSero and MLST all of which concerned phase variation. Since our findings indicated that MLST was not suited for identification of phase variation and SeqSero generally performed better in this regard, we rated the SeqSero result as more adequate. There was disagreement between SeqSero and MLST regarding 20 *S.* Typhimurium isolates of ST 34, which were classified as monophasic by MLST and biphasic by SeqSero. Since biphasic ST 34 isolates cannot be correctly classified by MLST we chose the SeqSero prediction for these cases. The same applied for monophasic ST 19 *S.* Typhimurium isolates, which were also not correctly classified by MLST. The two isolates, which carried a transposase in *fljB* (ERR2003330 and ERR2003327), were correctly predicted as monophasic by MLST and here we opted for the MLST prediction because we had already analyzed these isolates by mapping. In the 5 cases of prediction failure by SeqSero, we chose the MLST prediction as the serovar. This way, the percentage of correlation was increased to >99%. In summary the combination of both independent methods enabled the identification of potential misclassifications where a closer analysis was necessary and thus reduced the rate of error.

## Discussion

In this study we evaluated two genome-based *in silico* approaches and their combination for predicting *Salmonella* serotypes and their suitability for replacing classical serotyping. Table 3 summarizes the advantages and drawbacks of the three typing methods. We found that both tested prediction methods, the *in silico* serotyping approach by SeqSero 1.0 and the indirect serotype prediction with MLST yielded excellent correlation with our laboratory-based results analyzing 520 isolates from our strain collection (98% SeqSero, 95% MLST). Since our collection lacked a representative selection of strains of rare serotypes or higher subspecies we cannot rate the performance in this regard. Nonetheless it was representative of the most common human strains in Germany.

**Table 3.** Overview of advantages and drawbacks of the investigated typing methods and their sources of errors. Concerning classical serotyping we also referred to Hendriksen et al. 2009 ^13^.

| Typing Method | Advantage | Drawback | Main reasons for Errors | How to address sources of errors |
| --- | --- | --- | --- | --- |
| **Serotyping** | Directly determines phenotype | No typing of rough strains possible | Lack of experience with serotyping | Intensively trained staff |
|  | Well established method | Requires high quality antisera |  | Quality control mechanism |
| **SeqSero** | Classification analogous to classical serotyping | Genotype may not correspond to phenotype due to undetected mutations | Low sequence data quality | Quality control mechanism, e.g. of sequencing process |
|  | No assembly required | High quality sequencing data required (e.g. coverage, contamination) | Monophasic variants are only determined by lack of *fljB* | Improve detection method for monophasic variants |
|  | Can be automated |  |  |  |
| **MLST-based typing** | Provides phylogenetic information | High quality sequencing data required (e.g. coverage, contamination) | Low sequence data quality | Quality control mechanism, e.g. of sequencing process |
|  | Can be automated | Assembly recommended |  |  |

Our collection also included a novel serovar, derived from an outbreak related to sesame seeds ^18^. Interestingly, the antigenic formula of this novel serovar was correctly identified by SeqSero demonstrating its effectiveness for classifying novel serovars. Our correlation rate of 98% using raw reads matches very well the correlation rate determined by the developers of SeqSero of 98.7% using 308 CDC strains ^9^. However, the correlation rate found by Zhang and colleagues dropped to 92.6% when using a higher number of Isolates, i.e 3306 isolates from GenomeTrakr. Likewise, a recent study of 1041 environmental *Salmonella* isolates including a wider variety than our study yielded a correlation of 86% to classical serotyping ^15^. Recently, the developers of SeqSero presented a new version of the program named SeqSero 2.0 at the International Symposium on Salmonella and salmonellosis 2018 ^19^. SeqSero 2.0 can use SalmID in the assembly mode for subspecies identification of ambiguous serovars (www.github.com/hcdeenbakker/salmID). We did not test the assembly mode since it required the additional program SalmID, which we did not include in our assessment. We tested SeqSero 2.0 in its default k-mer based mode for reassessment of the ten cases where SeqSero failed. We found that with the default settings, SeqSero 2.0 also did not consistently detect monophasic variants of *S.* Typhimurium but showed improved performance in cases of high sequence variability.

Our results indicate that SeqSero does not reliably predict monophasic variants, in particular monophasic *S.* Typhimurium. Monophasic *S.* Typhimurium lacking *fljB* are correctly classified by SeqSero but atypical monophasic variants where *fljB* is present may be misclassified as biphasic. This is a potentially crucial limitation of the program as monophasic variants, especially of *S.* Typhimurium, are epidemiologically important and the latter comprise approximately 2/3 of the *S.* Typhimurium received at the NRC^20-24^. We suggest including the detection of additional factors to the *fljB* allele, which determine integrity of the second phase flagellar antigen. Also the algorithm for phase determination when using raw reads should be refined so that disruptions in the *fliC / fljB* genes can be detected in spite of the fact that the gene is fully present.

Regarding MLST, it was foreseeable by examining the strains in Enterobase that a clear classification between monophasic and biphasic *S.* Typhimurium based on ST would not be possible. Achtman et al. did not find a correlation between ST and monophasic *S.* Typhimurium when they analyzed a large and diverse collection ^4^. On the other hand, it was reported that Italian and UK monophasic *S.* Typhimurium strains belonged to ST 34 ^20,21^. Petrovska et al. showed that the current monophasic epidemic *S.* Typhimurium strains evolved from at least three independent events ^21^. The monophasic strains of our collection predominantly belong to the current European ST 34 epidemic clone, therefore a good correlation for monophasic strains of ST 34 was obtained with MLST. On the other hand, biphasic strains of ST 34 were misclassified as monophasic. We therefore conclude that the classical MLST scheme alone is not able to clearly distinguish between monophasic and biphasic *S.* Typhimurium due to their polyphyletic nature. Our results further indicate that clustering by core genome MLST does also not improve classification according to flagellin expression. Since recent studies have found *S.* Typhimurium regions, which seem characteristic for certain monophasic variants it may be possible to develop an additional scheme based on the presence/absence of such specific genes to reliably identify monophasic variants of *S.* Typhimurium ^21,24^.

We obtained the highest correlation to classical serotyping when we combined the predictions of SeqSero and MLST because the two methods use independent approaches for serotype determination and thereby complemented each other. Since SeqSero directly generates an antigenic formula, we rated its output as more adequate than the indirect determination by MLST. Nonetheless, with the additional information provided by MLST, it was possible to clarify all ambiguous predictions by SeqSero because the serovars, which shared the same antigenic formula, had different STs. Our results also indicate that MLST might even perform better in classifying rough strains than SeqSero. The combined prediction increased robustness because miscorrelating predictions of the two programs gave rise to more detailed analysis. Currently there are two tools available, which use the combined prediction of *in silico* serotyping and MLST. One is SalmonellaTypeFinder, which uses SeqSero and MLST and thus has the potential of performing well (https://bitbucket.org/genomicepidemiology/salmonellatypefinder/src/master/). We did not evaluate this tool in our study because it uses a different MLST calling algorithm than we routinely do and has not been published yet. The second tool is SISTR, which predicts the serotype with the help of *in silico* genoserotyping and validates the results with core genome MLST ^10^. We did not evaluate SISTR because it requires assembled genomes. However, it performed very well in a previous report^14^. A combination of genome-based serotyping and MLST is also advocated by other governmental agencies like Public Health England, who use MOST and Public Health Agency of Canada who use SISTR ^14^.

## Conclusion

SeqSero is an *in silico* serotyping tool generating an antigenic formula directly comparable to classical serotyping. MLST provides important phylogenetic information and is able to distinguish serovars with the same antigenic formula. The concomitant use of both tools seems best suited for *in silico* strain characterization to obtain the utmost information and a robust prediction. Nevertheless, some improvements are necessary to differentiate monophasic from biphasic strains. If the serotype is predicted by these two independent methods, a disagreement could indicate a potential problem requiring further investigation. Since we obtained a correlation rate of >99% for SeqSero in combination with MLST, we conclude that the here investigated *in silico* typing tools could in combination outperform the current gold standard of phenotypic serotyping and could become the new gold standard.

## Methods

### Short read sequencing

Whole genome sequencing was performed at the NRC or at the Robert Koch-Institute’s sequencing core facility on a MiSeq benchtop sequencer using Illumina’s MiSeq Reagent Kit v3, yielding 2 x 300 bp paired end reads. Adapter-clipped reads were obtained from the sequencing unit and used in this study without additional processing unless stated otherwise. Sequencing was repeated for cases not meeting the minimal read number of 100,000 (Fig. S1). The fastq files of the paired-end sequence reads are available from the European Nucleotide Archive under the project numbers PRJEB30317 & PRJEB16326. Project PRJEB16326 is part of EU COMPARE (https://www.compare-europe.eu/) and a subset of the German samples of that project have been included in this study.

### SeqSero 1.0/SeqSero 2.0

The SeqSero 1.0 command line tool was downloaded from Github (https://github.com/denglab/SeqSero) and an official Debian package was created, which is available from https://blends.debian.org/med/tasks/bio. The installed program was then embedded into a script for batch analysis. Illumina MiSeq paired-end reads were directly used for serotype prediction. Apart from choosing the correct mode for the input data, i.e. single-end, paired-end, interleaved or assembled, the program offers no additional options. During drafting of this manuscript an alpha test version of SeqSero 2.0 became available from Github (https://github.com/denglab/SeqSero 2.0). We used SeqSero 2.0 with its default setting (k-mer based mode) only to analyze isolates where SeqSero 1.0 did not produce a correlating result to classical serotyping.

### Ridom SeqSphere settings and allele calling procedure

For MLST analysis we used the 7 gene MLST scheme from Achtman *et al.* embedded in the Ridom SeqSphere^+^ software (Ridom GmbH, Münster, Germany) ^4^. Please note, that in spite of the fact that the scheme recommends *de novo* assembly of raw reads, we used mapping in order to save time and resources. Using the raw reads, the pipeline quality-trimmed and mapped the Illumina MiSeq reads against the reference genome *S.* Typhimurium LT2 (GenBank AE006468.2) using the build-in Burrows-Wheler Aligner in the default mode. This ideally yielded allele numbers for the seven housekeeping genes and the corresponding sequence type (ST). If Ridom SeqSphere^+^ was not able to assign a ST there were generally two reasons: either low data quality (‘Target QC procedure failure’) or it was a potential new ST. For cases of low sequence quality sequencing was repeated (Fig. S1). For phylogenetic analysis of monophasic and biphasic *S.* Typhimurium isolates the Enterobase core genome MLST scheme was used in SeqSphere^+^.

### Assigning Sequence types and corresponding serotypes with Enterobase

The obtained MLST sequence types were entered into Enterobase to find corresponding serotypes from the database and if available the e-burst groups (eBGs). eBGs determination is based on an algorithm, which identifies the relationship of isolates with similar genotypes^17^. Enterobase periodically confers official eBG numbers to new eBGs.

If Ridom SeqSphere^+^ reported a potential new ST we uploaded the NGS data of the respective isolates to Enterobase in order to obtain an official ST.

### De novo assembly and mapping

For isolates with non-correlating results *de novo* assembly was performed using A5 or SPAdes ^25,26^ Some isolates where further analyzed by mapping the raw reads against specific loci using the Geneious mapper or Bowtie2 in Geneious (www.geneious.com).

## Data Availability

The raw sequence reads analyzed in this study are publicly available at the European Nucleotide Archive under the project accession numbers PRJEB30317 and PRJEB16326. PRJEB16326 is part of COMPARE and a subset of the German samples has been included in the current study. An overview of all strains and metadata is given in Table S1.

## Acknowledgements

We thank Monique Duwe, Marita Wahnfried and Susanne Kulbe for excellent technical assistance. We thank the RKI NGS sequencing team in Berlin and Wernigerode. We are also thankful to all laboratories that sent strains to the NRC. The authors acknowledge funding by the EU COMPARE project within the Horizon 2020 (AF).

## Author contributions

SB performed the *in silico* analysis and drafted the manuscript. SS supervised the serotyping and sequencing, advised in cgMLST analysis and provided input to the manuscript. AT wrote scripts for batch analysis and extraction of results. AF conceived and supervised the project and provided input to the manuscript.

## Competing interests

The authors declare no competing interests.

