## supplementary data for "Can genome-based analysis be the new gold-standard for routine Salmonella serotyping?"

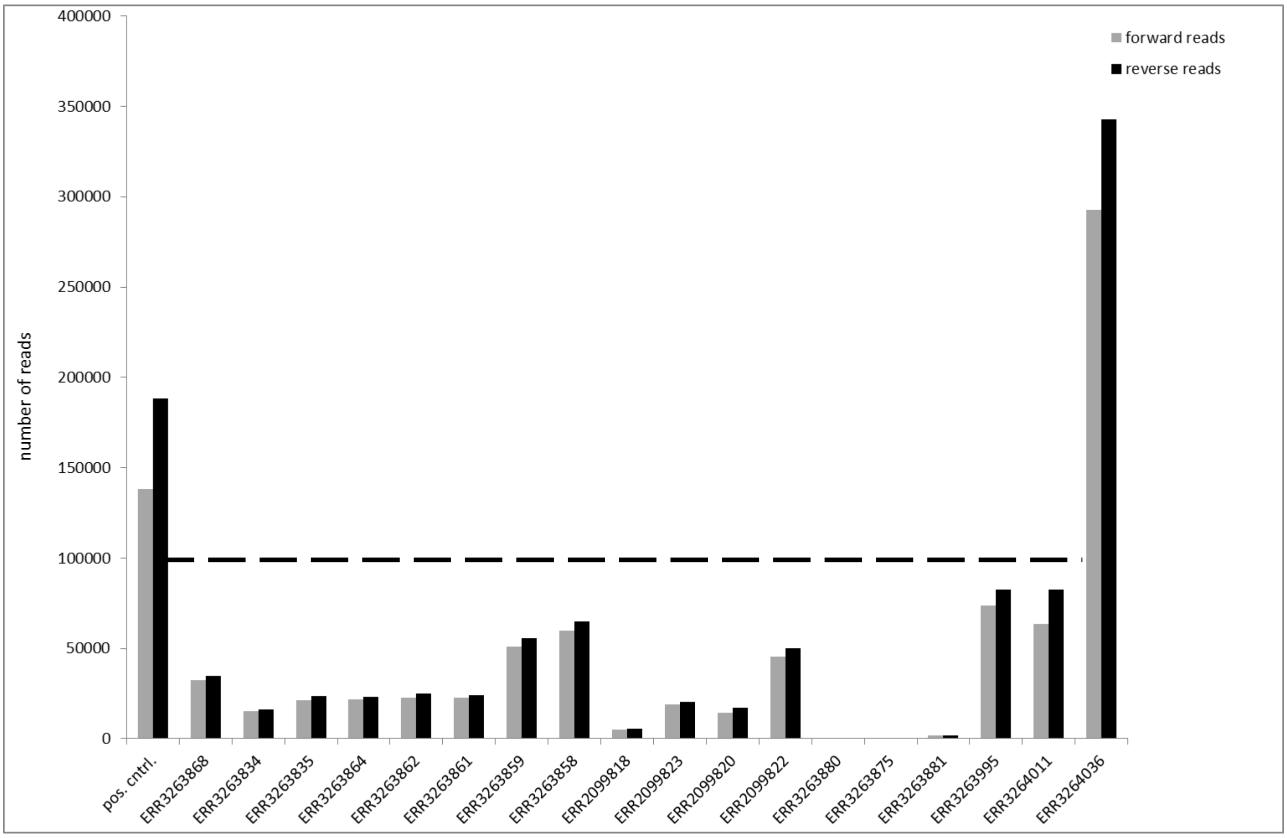


Fig. S1. Overview of data quality of sequences with failed serotype prediction by SeqSero. Data quality is represented by the number of reads in the forward and reverse read files of paired-end reads. Isolates with unsuccessful serotype prediction by SeqSero are shown as well as a positive control (ERR3263813) with successful serotype prediction. Please note that for isolate ERR3264036 serotype prediction failed in spite of sufficient data quality because of missing O-7 antigen locus (Fig. S3). The dashed line indicates the threshold of 100,000 reads per direction for paired end reads to achieve a theoretical coverage of 10-fold.

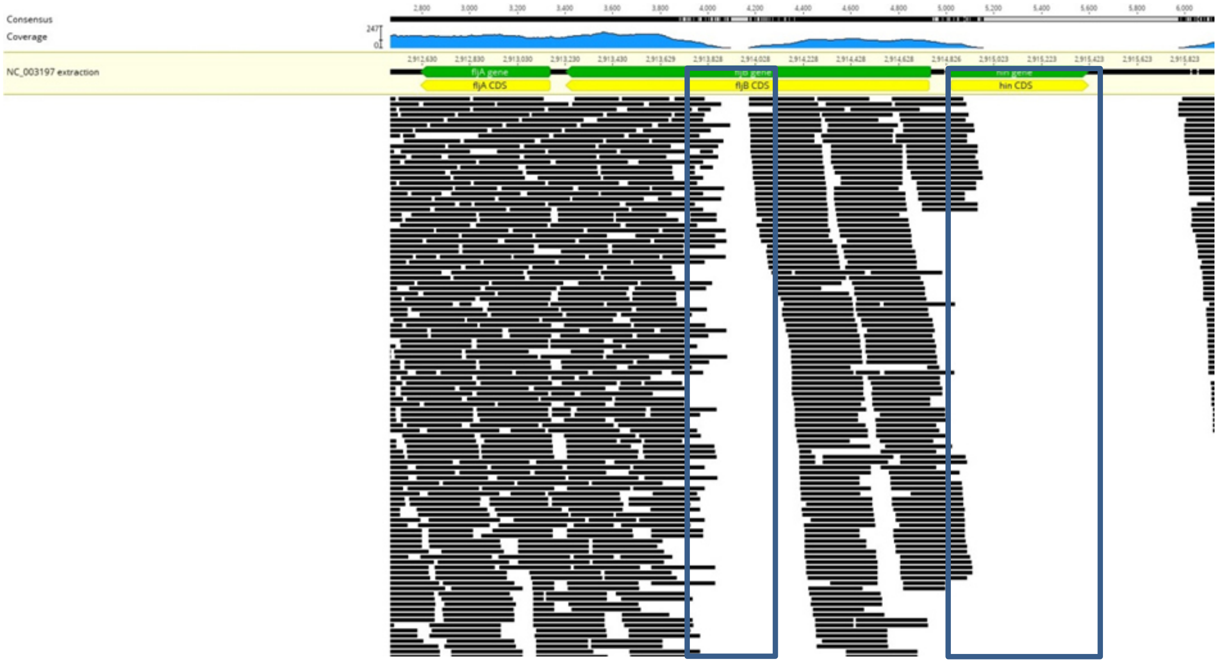


Fig. S2. Isolate ERR2003330 lacks the invertase gene *hin* as well as the central part of the *fljB* gene. Raw reads of Isolate ERR2003330 were mapped against the conserved *fljAB* region extracted from *S.* Typhimurium LT2 using the Geneious mapper in Geneious.

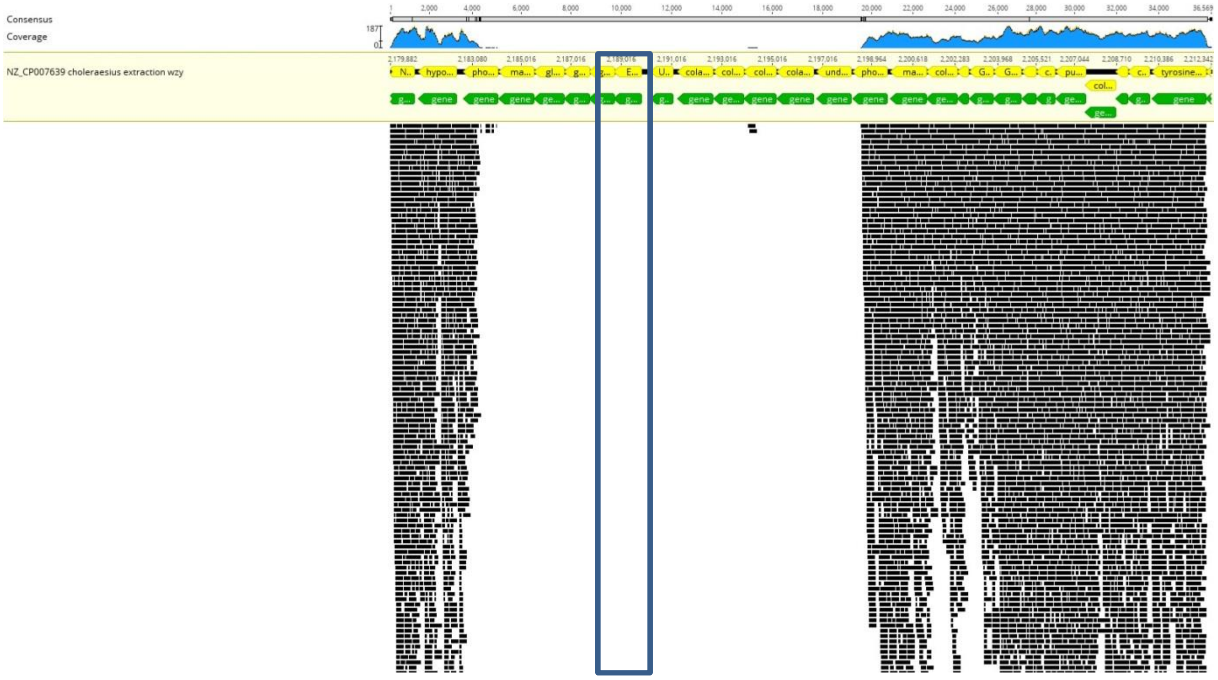


Fig. S3. *S.* Choleraesuis O-7 antigen locus and surrounding region are missing in isolate ERR3264036. Raw reads of isolate ERR3264036 were mapped against region EL48_RS10955- EL48_RS11010, including the *wzy* *l*ocus EL48_RS10980 of *S.* Choleraesuis NZ_CP007639 using Bowtie2 in Geneious.

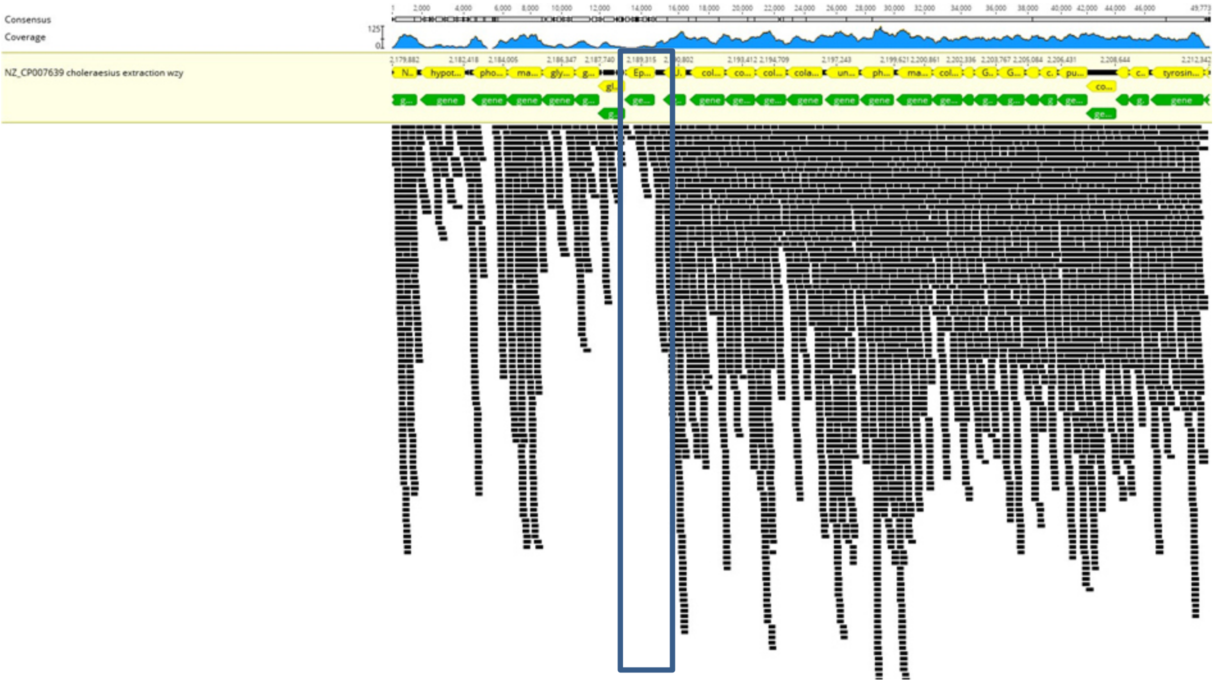


Fig. S4. Poor read coverage of *S.* Choleraesuis O-7 antigen locus in isolate ERR3264063. Raw reads of isolate ERR3264063 were mapped against region EL48_RS10955- EL48_RS11010, including the *wzy* *l*ocus EL48_RS10980 of *S.* Choleraesuis NZ_CP007639 using Bowtie2 in Geneious. This figure is also representative for isolates ERR3264067 and ERR3264066.

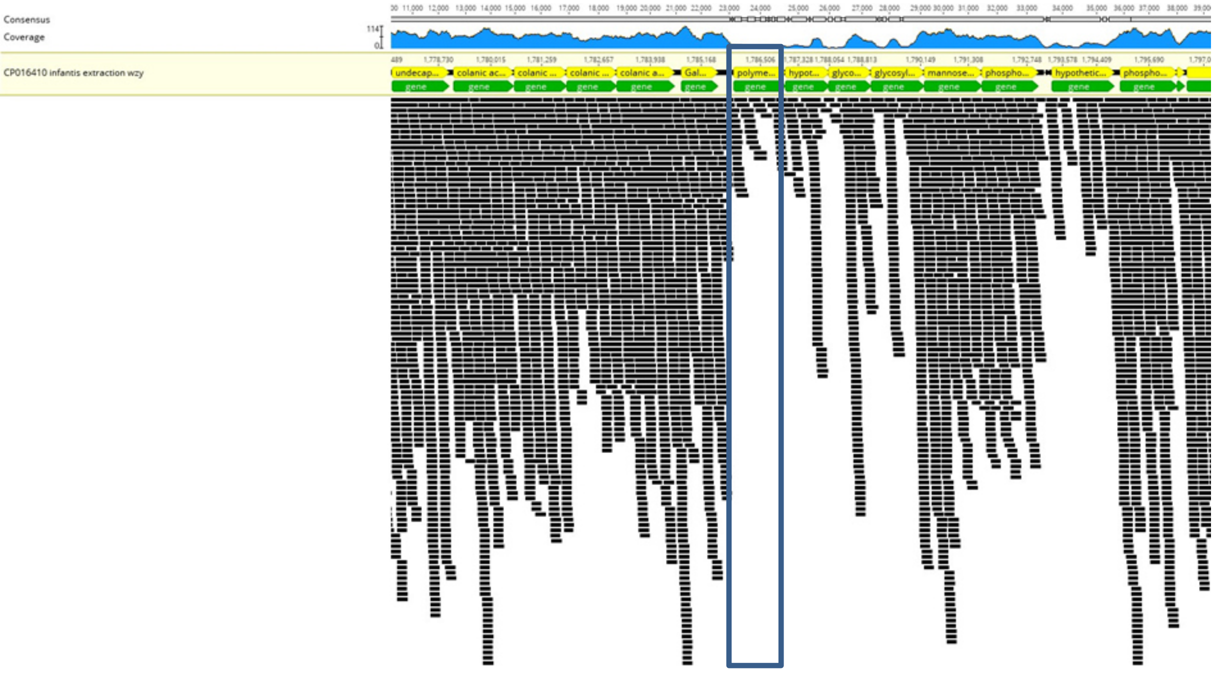


Fig. S5. Poor read coverage of *S. Infantis* O-7 antigen locus in isolate ERR3264076. Raw reads of isolate ERR3264076 were mapped against region APL92_08575- APL92_08690, including the *wzy* *l*ocus APL92_08605 of *S.* Infantis CP016410 using Bowtie2 in Geneious.

Table S1. Overview of NRC dataset for the study Banerji et al.: 'Can genome-based analysis be the new gold-standard for routine Salmonella serotyping?'. The dataset consists of 520 whole genome sequences, which have been submitted to ENA.

| ENA run number | RKI number | Source | Year of Isolation | Country of Isolation | Serovar | ENA Project |
|---|---|---|---|---|---|---|
| ERR3263618 | RKI_09-07379 | Human | 2009 | Germany | S. Infantis | PRJEB30317 |
| ERR3263619 | RKI_09-07426 | Human | 2009 | Germany | S. Infantis | PRJEB30317 |
| ERR3263620 | RKI_09-07751 | Human | 2009 | Germany | S. Infantis | PRJEB30317 |
| ERR3263621 | RKI_01-11525 | Human | 2001 | Germany | S. Sundsvall | PRJEB30317 |
| ERR3263622 | RKI_04-05621 | Meat | 2004 | Germany | S. Derby | PRJEB30317 |
| ERR3263623 | RKI_06-01900 | Human | 2006 | Germany | S. Typhimurium mono | PRJEB30317 |
| ERR3263624 | RKI_07-00493 | Human | 2007 | Germany | S. Kintambo | PRJEB30317 |
| ERR3263625 | RKI_07-01167 | Human | 2007 | Germany | S. Derby | PRJEB30317 |
| ERR3263626 | RKI_07-01409 | Human | 2007 | Germany | S. Poano | PRJEB30317 |
| ERR3263627 | RKI_07-02601 | Human | 2007 | Germany | S. Derby | PRJEB30317 |
| ERR3263628 | RKI_07-04162 | Human | 2007 | Germany | S. Derby | PRJEB30317 |
| ERR3263629 | RKI_07-05911 | Human | 2007 | Germany | S. Derby | PRJEB30317 |
| ERR3263630 | RKI_08-01395 | Human | 2008 | Germany | S. Derby | PRJEB30317 |
| ERR3263631 | RKI_09-06166 | Human | 2009 | Germany | S. Kintambo | PRJEB30317 |
| ERR3263632 | RKI_11-01214 | Human | 2011 | Germany | S. Kintambo | PRJEB30317 |
| ERR3263633 | RKI_11-01215 | Human | 2011 | Germany | S. Poano | PRJEB30317 |
| ERR3263634 | RKI_12-00623 | Meat | 2012 | Germany | S. Derby | PRJEB30317 |
| ERR3263635 | RKI_12-01254 | Human | 2012 | Germany | S. Derby | PRJEB30317 |
| ERR3263636 | RKI_12-01969 | Human | 2012 | Germany | S. Derby | PRJEB30317 |
| ERR3263637 | RKI_12-03084 | Human | 2012 | Germany | S. Derby | PRJEB30317 |
| ERR3263638 | RKI_12-05564 | Poultry | 2012 | Germany | S. Derby | PRJEB30317 |
| ERR3263639 | RKI_12-06007 | Human | 2012 | Germany | S. Derby | PRJEB30317 |
| ERR3263640 | RKI_13-01828 | Human | 2013 | Germany | S. Infantis | PRJEB30317 |
| ERR3263641 | RKI_13-02310 | Human | 2013 | Germany | S. Infantis | PRJEB30317 |
| ERR3263642 | RKI_13-02356 | Cattle | 2013 | Germany | S. Infantis | PRJEB30317 |
| ERR3263643 | RKI_13-02889 | Human | 2013 | Germany | S. Derby | PRJEB30317 |
| ERR3263644 | RKI_13-03088 | Human | 2013 | Germany | S. Muenchen | PRJEB30317 |
| ERR3263645 | RKI_13-03099 | Human | 2013 | Germany | S. Muenchen | PRJEB30317 |
| ERR3263646 | RKI_13-03108 | Meat | 2013 | Germany | S. Muenchen | PRJEB30317 |
| ERR3263647 | RKI_13-03192 | Human | 2013 | Germany | S. Muenchen | PRJEB30317 |
| ERR3263648 | RKI_13-03193 | Human | 2013 | Germany | S. Muenchen | PRJEB30317 |
| ERR3263649 | RKI_13-03387 | Pig | 2013 | Germany | S. Muenchen | PRJEB30317 |
| ERR3263650 | RKI_13-03389 | Pig | 2013 | Germany | S. Muenchen | PRJEB30317 |
| ERR3263651 | RKI_13-04497 | Human | 2013 | Germany | S. Derby | PRJEB30317 |
| ERR3263652 | RKI_13-04650 | Human | 2013 | Germany | S. Derby | PRJEB30317 |
| ERR3263653 | RKI_13-05246 | Human | 2013 | Germany | S. Derby | PRJEB30317 |
| ERR3263654 | RKI_13-05853 | Human | 2013 | Germany | S. Derby | PRJEB30317 |
| ERR3263655 | RKI_13-05979 | Human | 2013 | Germany | S. Derby | PRJEB30317 |
| ERR3263656 | RKI_14-00018 | Human | 2014 | Germany | S. Derby | PRJEB30317 |
| ERR3263657 | RKI_14-00019 | Human | 2014 | Germany | S. Derby | PRJEB30317 |
| ERR3263658 | RKI_14-00111 | Human | 2014 | Germany | S. Derby | PRJEB30317 |
| ERR3263659 | RKI_14-00117 | Human | 2014 | Germany | S. Derby | PRJEB30317 |
| ERR3263660 | RKI_14-00150 | Human | 2014 | Germany | S. Derby | PRJEB30317 |
| ERR3263661 | RKI_14-00162 | Human | 2014 | Germany | S. Derby | PRJEB30317 |
| ERR3263662 | RKI_14-00171 | Human | 2014 | Germany | S. Derby | PRJEB30317 |
| ERR3263663 | RKI_14-00238 | Human | 2014 | Germany | S. Derby | PRJEB30317 |
| ERR3263664 | RKI_14-00239 | Human | 2014 | Germany | S. Derby | PRJEB30317 |
| ERR3263665 | RKI_14-00383 | Human | 2014 | Germany | S. Derby | PRJEB30317 |
| ERR3263666 | RKI_14-00404 | Human | 2014 | Germany | S. Derby | PRJEB30317 |
| ERR3263667 | RKI_14-00426 | Human | 2014 | Germany | S. Derby | PRJEB30317 |
| ERR3263668 | RKI_14-00470 | Human | 2014 | Germany | S. Derby | PRJEB30317 |
| ERR3263669 | RKI_14-00515 | Human | 2014 | Germany | S. Derby | PRJEB30317 |
| ERR3263670 | RKI_14-00537 | Human | 2014 | Germany | S. Muenchen | PRJEB30317 |
| ERR3263671 | RKI_14-00626 | Human | 2014 | Germany | S. Derby | PRJEB30317 |
| ERR3263672 | RKI_14-00651 | Human | 2014 | Germany | S. Derby | PRJEB30317 |
| ERR3263673 | RKI_14-00672 | Human | 2014 | Germany | S. Derby | PRJEB30317 |
| ERR3263674 | RKI_14-00683 | Human | 2014 | Germany | S. Derby | PRJEB30317 |
| ERR3263675 | RKI_14-00730 | Human | 2014 | Germany | S. Derby | PRJEB30317 |
| ERR3263676 | RKI_14-00731 | Human | 2014 | Germany | S. Derby | PRJEB30317 |
| ERR3263677 | RKI_14-00748 | Human | 2014 | Germany | S. Derby | PRJEB30317 |
| ERR3263678 | RKI_14-00751 | Human | 2014 | Germany | S. Derby | PRJEB30317 |
| ERR3263679 | RKI_14-00758 | Human | 2014 | Germany | S. Derby | PRJEB30317 |
| ERR3263680 | RKI_14-00841 | Human | 2014 | Germany | S. Muenchen | PRJEB30317 |
| ERR3263681 | RKI_14-00860 | Human | 2014 | Germany | S. Derby | PRJEB30317 |
| ERR3263682 | RKI_14-00922 | Human | 2014 | Germany | S. Derby | PRJEB30317 |
| ERR3263683 | RKI_14-01037 | Human | 2014 | Germany | S. Derby | PRJEB30317 |
| ERR3263684 | RKI_14-01040 | Human | 2014 | Germany | S. Derby | PRJEB30317 |
| ERR3263685 | RKI_14-01041 | Human | 2014 | Germany | S. Derby | PRJEB30317 |
| ERR3263686 | RKI_14-01066 | Human | 2014 | Germany | S. Derby | PRJEB30317 |
| ERR3263687 | RKI_14-01163 | Human | 2014 | Germany | S. Derby | PRJEB30317 |
| ERR3263688 | RKI_14-01398 | Human | 2014 | Germany | S. Derby | PRJEB30317 |
| ERR3263689 | RKI_14-01773 | Human | 2014 | Germany | S. Derby | PRJEB30317 |
| ERR3263690 | RKI_14-02487 | Human | 2014 | Germany | S. Muenchen | PRJEB30317 |

| | | | | | | |
|---|---|---|---|---|---|---|
| ERR3263691 | RKI_14-02489 | Human | 2014 | Germany | S. Muenchen | PRJEB30317 |
| ERR3263692 | RKI_14-02527 | Human | 2014 | Germany | S. Muenchen | PRJEB30317 |
| ERR3263693 | RKI_14-02618 | Human | 2014 | Germany | S. Muenchen | PRJEB30317 |
| ERR3263694 | RKI_14-02626 | Human | 2014 | Germany | S. Muenchen | PRJEB30317 |
| ERR3263695 | RKI_14-02631 | Human | 2014 | Germany | S. Muenchen | PRJEB30317 |
| ERR3263696 | RKI_14-02632 | Human | 2014 | Germany | S. Muenchen | PRJEB30317 |
| ERR3263697 | RKI_14-02676 | Human | 2014 | Germany | S. Muenchen | PRJEB30317 |
| ERR3263698 | RKI_14-02681 | Meat | 2014 | Germany | S. Muenchen | PRJEB30317 |
| ERR3263699 | RKI_14-02705 | Human | 2014 | Germany | S. Muenchen | PRJEB30317 |
| ERR3263700 | RKI_14-02717 | Meat | 2014 | Germany | S. Muenchen | PRJEB30317 |
| ERR3263701 | RKI_14-02718 | Meat | 2014 | Germany | S. Muenchen | PRJEB30317 |
| ERR3263702 | RKI_14-02719 | Human | 2014 | Germany | S. Muenchen | PRJEB30317 |
| ERR3263703 | RKI_14-03337 | Pig | 2014 | Germany | S. Muenchen | PRJEB30317 |
| ERR3263704 | RKI_14-03339 | Pig | 2014 | Germany | S. Muenchen | PRJEB30317 |
| ERR3263705 | RKI_14-03538 | Human | 2014 | Germany | S. Muenchen | PRJEB30317 |
| ERR3263706 | RKI_14-03731 | Human | 2014 | Germany | S. Infantis | PRJEB30317 |
| ERR3263707 | RKI_14-04275 | Human | 2014 | Germany | S. Enteritidis | PRJEB30317 |
| ERR3263708 | RKI_14-04296 | Human | 2014 | Germany | S. Enteritidis | PRJEB30317 |
| ERR3263709 | RKI_14-04310 | Human | 2014 | Germany | S. Enteritidis | PRJEB30317 |
| ERR3263710 | RKI_14-04552 | Human | 2014 | Germany | S. Enteritidis | PRJEB30317 |
| ERR3263711 | RKI_14-04639 | Human | 2014 | Germany | S. Enteritidis | PRJEB30317 |
| ERR3263712 | RKI_14-04870 | Human | 2014 | Germany | S. Enteritidis | PRJEB30317 |
| ERR3263713 | RKI_14-05567 | Human | 2014 | Germany | S. Enteritidis | PRJEB30317 |
| ERR3263714 | RKI_14-05569 | Human | 2014 | Germany | S. Enteritidis | PRJEB30317 |
| ERR3263715 | RKI_14-05795 | Human | 2014 | Germany | S. Enteritidis | PRJEB30317 |
| ERR3263716 | RKI_14-05946 | Human | 2014 | Germany | S. Enteritidis | PRJEB30317 |
| ERR3263717 | RKI_14-06012 | Human | 2014 | Germany | S. Enteritidis | PRJEB30317 |
| ERR3263718 | RKI_14-06145 | Human | 2014 | Germany | S. Enteritidis | PRJEB30317 |
| ERR3263719 | RKI_14-06148 | Human | 2014 | Germany | S. Enteritidis | PRJEB30317 |
| ERR3263720 | RKI_14-06175 | Human | 2014 | Germany | S. Enteritidis | PRJEB30317 |
| ERR3263721 | RKI_14-06388 | Human | 2014 | Germany | S. Enteritidis | PRJEB30317 |
| ERR3263722 | RKI_15-00251 | Human | 2015 | Germany | S. Typhi | PRJEB30317 |
| ERR3263723 | RKI_15-00325 | Human | 2015 | Germany | S. Enteritidis | PRJEB30317 |
| ERR3263724 | RKI_15-00342 | Human | 2015 | Germany | S. Typhi | PRJEB30317 |
| ERR3263725 | RKI_15-00347 | Human | 2015 | Germany | S. Enteritidis | PRJEB30317 |
| ERR3263726 | RKI_15-00348 | Human | 2015 | Germany | S. Enteritidis | PRJEB30317 |
| ERR3263727 | RKI_15-00349 | Human | 2015 | Germany | S. Enteritidis | PRJEB30317 |
| ERR3263728 | RKI_15-00350 | Human | 2015 | Germany | S. Enteritidis | PRJEB30317 |
| ERR3263729 | RKI_15-00355 | Human | 2015 | Germany | S. Typhi | PRJEB30317 |
| ERR3263730 | RKI_15-00407 | Human | 2015 | Germany | S. Typhi | PRJEB30317 |
| ERR3263731 | RKI_15-00807 | Human | 2015 | Germany | S. Enteritidis | PRJEB30317 |
| ERR3263732 | RKI_15-01127 | Human | 2015 | Germany | S. Typhi | PRJEB30317 |
| ERR3263733 | RKI_15-01202 | Human | 2015 | Germany | S. Typhi | PRJEB30317 |
| ERR3263734 | RKI_15-01240 | Human | 2015 | Germany | S. Typhi | PRJEB30317 |
| ERR3263735 | RKI_15-01243 | Human | 2015 | Germany | S. Typhi | PRJEB30317 |
| ERR3263736 | RKI_15-01252 | Human | 2015 | Germany | S. Typhi | PRJEB30317 |
| ERR3263737 | RKI_15-01462 | Human | 2015 | Germany | S. Typhi | PRJEB30317 |
| ERR3263738 | RKI_15-01478 | Not Provide | 2015 | Germany | S. Typhi | PRJEB30317 |
| ERR3263739 | RKI_15-01485 | Human | 2015 | Germany | S. Typhi | PRJEB30317 |
| ERR3263740 | RKI_15-01563 | Human | 2015 | Germany | S. Enteritidis | PRJEB30317 |
| ERR3263741 | RKI_15-01620 | Human | 2015 | Germany | S. Typhi | PRJEB30317 |
| ERR3263742 | RKI_15-01837 | Human | 2015 | Germany | S. Typhi | PRJEB30317 |
| ERR3263743 | RKI_15-01874 | Not Provide | 2015 | Germany | S. Typhi | PRJEB30317 |
| ERR3263744 | RKI_15-02000 | Human | 2015 | Germany | S. Typhi | PRJEB30317 |
| ERR3263745 | RKI_15-02628 | Human | 2015 | Germany | S. Typhi | PRJEB30317 |
| ERR3263746 | RKI_15-02720 | Human | 2015 | Germany | S. Typhi | PRJEB30317 |
| ERR3263747 | RKI_15-02721 | Human | 2015 | Germany | S. Typhi | PRJEB30317 |
| ERR3263748 | RKI_15-02844 | Human | 2015 | Germany | S. Enteritidis | PRJEB30317 |
| ERR3263749 | RKI_15-02845 | Human | 2015 | Germany | S. Enteritidis | PRJEB30317 |
| ERR3263750 | RKI_15-02846 | Human | 2015 | Germany | S. Enteritidis | PRJEB30317 |
| ERR3263751 | RKI_15-02949 | Human | 2015 | Germany | S. Enteritidis | PRJEB30317 |
| ERR3263752 | RKI_15-03059 | Human | 2015 | Germany | S. Enteritidis | PRJEB30317 |
| ERR3263753 | RKI_15-03112 | Human | 2015 | Germany | S. Enteritidis | PRJEB30317 |
| ERR3263754 | RKI_15-03113 | Human | 2015 | Germany | S. Enteritidis | PRJEB30317 |
| ERR3263755 | RKI_15-03114 | Human | 2015 | Germany | S. Typhi | PRJEB30317 |
| ERR3263756 | RKI_15-03208 | Human | 2015 | Germany | S. Enteritidis | PRJEB30317 |
| ERR3263757 | RKI_15-03215 | Human | 2015 | Germany | S. Stourbridge | PRJEB30317 |
| ERR3263758 | RKI_15-03226 | Human | 2015 | Germany | S. Enteritidis | PRJEB30317 |
| ERR3263759 | RKI_15-03227 | Human | 2015 | Germany | S. Enteritidis | PRJEB30317 |
| ERR3263760 | RKI_15-03228 | Human | 2015 | Germany | S. Enteritidis | PRJEB30317 |
| ERR3263761 | RKI_15-03229 | Human | 2015 | Germany | S. Enteritidis | PRJEB30317 |
| ERR3263762 | RKI_15-03258 | Human | 2015 | Germany | S. Typhi | PRJEB30317 |
| ERR3263763 | RKI_15-03274 | Human | 2015 | Germany | S. Enteritidis | PRJEB30317 |
| ERR3263764 | RKI_15-03305 | Human | 2015 | Germany | S. Typhi | PRJEB30317 |
| ERR3263765 | RKI_15-03335 | Human | 2015 | Germany | S. Enteritidis | PRJEB30317 |
| ERR3263766 | RKI_15-03367 | Human | 2015 | Germany | S. Enteritidis | PRJEB30317 |

| | | | | | | |
|---|---|---|---|---|---|---|
| ERR3263767 | RKI_15-03370 | Human | 2015 | Germany | S. Enteritidis | PRJEB30317 |
| ERR3263768 | RKI_15-03373 | Human | 2015 | Germany | S. Enteritidis | PRJEB30317 |
| ERR3263769 | RKI_15-03374 | Human | 2015 | Germany | S. Enteritidis | PRJEB30317 |
| ERR3263770 | RKI_15-03375 | Human | 2015 | Germany | S. Enteritidis | PRJEB30317 |
| ERR3263771 | RKI_15-03440 | Human | 2015 | Germany | S. Enteritidis | PRJEB30317 |
| ERR3263772 | RKI_15-03455 | Human | 2015 | Germany | S. Enteritidis | PRJEB30317 |
| ERR3263773 | RKI_15-03456 | Human | 2015 | Germany | S. Enteritidis | PRJEB30317 |
| ERR3263774 | RKI_15-03486 | Human | 2015 | Germany | S. Enteritidis | PRJEB30317 |
| ERR3263775 | RKI_15-03487 | Human | 2015 | Germany | S. Enteritidis | PRJEB30317 |
| ERR3263776 | RKI_15-03489 | Human | 2015 | Germany | S. Enteritidis | PRJEB30317 |
| ERR3263777 | RKI_15-03538 | Human | 2015 | Germany | S. Enteritidis | PRJEB30317 |
| ERR3263778 | RKI_15-03552 | Human | 2015 | Germany | S. Enteritidis | PRJEB30317 |
| ERR3263779 | RKI_15-03579 | Human | 2015 | Germany | S. Enteritidis | PRJEB30317 |
| ERR3263780 | RKI_15-03582 | Poultry | 2015 | Germany | S. Enteritidis | PRJEB30317 |
| ERR3263781 | RKI_15-03584 | Poultry | 2015 | Germany | S. Enteritidis | PRJEB30317 |
| ERR3263782 | RKI_15-03614 | Human | 2015 | Germany | S. Enteritidis | PRJEB30317 |
| ERR3263783 | RKI_15-03637 | Layer Eggs | 2015 | Germany | S. Enteritidis | PRJEB30317 |
| ERR3263784 | RKI_15-03654 | Human | 2015 | Germany | S. Enteritidis | PRJEB30317 |
| ERR3263785 | RKI_15-03747 | Human | 2015 | Germany | S. Typhi | PRJEB30317 |
| ERR3263786 | RKI_15-03785 | Human | 2015 | Germany | S. Paratyphi B | PRJEB30317 |
| ERR3263787 | RKI_15-03791 | Human | 2015 | Germany | S. Typhi | PRJEB30317 |
| ERR3263788 | RKI_15-03832 | Human | 2015 | Germany | S. Enteritidis | PRJEB30317 |
| ERR3263789 | RKI_15-03840 | Human | 2015 | Germany | S. Paratyphi B | PRJEB30317 |
| ERR3263790 | RKI_15-03906 | Human | 2015 | Germany | S. Typhi | PRJEB30317 |
| ERR3263791 | RKI_15-03919 | Human | 2015 | Germany | S. Typhi | PRJEB30317 |
| ERR3263792 | RKI_15-04027 | Human | 2015 | Germany | S. Typhi | PRJEB30317 |
| ERR3263793 | RKI_15-04043 | Human | 2015 | Germany | S. Infantis | PRJEB30317 |
| ERR3263794 | RKI_15-04049 | Human | 2015 | Germany | S. Paratyphi B | PRJEB30317 |
| ERR3263795 | RKI_15-04101 | Human | 2015 | Germany | S. Typhi | PRJEB30317 |
| ERR3263796 | RKI_15-04114 | Human | 2015 | Germany | S. Typhi | PRJEB30317 |
| ERR3263797 | RKI_15-04165 | Human | 2015 | Germany | S. Typhi | PRJEB30317 |
| ERR3263798 | RKI_15-04264 | Human | 2015 | Germany | S. Typhi | PRJEB30317 |
| ERR3263799 | RKI_15-04397 | Human | 2015 | Germany | S. Typhi | PRJEB30317 |
| ERR3263800 | RKI_15-04398 | Human | 2015 | Germany | S. Typhi | PRJEB30317 |
| ERR3263801 | RKI_15-04481 | Human | 2015 | Germany | S. Typhi | PRJEB30317 |
| ERR3263802 | RKI_15-04494 | Human | 2015 | Germany | S. Typhi | PRJEB30317 |
| ERR3263803 | RKI_15-04495 | Human | 2015 | Germany | S. Typhi | PRJEB30317 |
| ERR3263804 | RKI_15-04496 | Human | 2015 | Germany | S. Typhi | PRJEB30317 |
| ERR3263805 | RKI_15-04497 | Human | 2015 | Germany | S. Typhi | PRJEB30317 |
| ERR3263806 | RKI_15-04510 | Human | 2015 | Germany | S. Stourbridge | PRJEB30317 |
| ERR3263807 | RKI_15-04638 | Human | 2015 | Germany | S. Typhi | PRJEB30317 |
| ERR3263808 | RKI_15-04657 | Human | 2015 | Germany | S. Typhi | PRJEB30317 |
| ERR3263809 | RKI_15-04702 | Human | 2015 | Germany | S. Enteritidis | PRJEB30317 |
| ERR3263810 | RKI_15-01116 | Human | 2015 | Germany | S. Infantis | PRJEB30317 |
| ERR3263811 | RKI_15-02397 | Human | 2015 | Germany | S. Infantis | PRJEB30317 |
| ERR3263812 | RKI_15-03230 | Human | 2015 | Germany | S. Infantis | PRJEB30317 |
| ERR3263813 | RKI_16-00051 | Human | 2016 | Germany | S. Typhi | PRJEB30317 |
| ERR3263814 | RKI_18-05968 | Human | 2018 | Germany | S. Typhimurium | PRJEB30317 |
| ERR3263815 | RKI_16-00056 | Human | 2016 | Germany | S. Mbandaka | PRJEB30317 |
| ERR3263816 | RKI_16-00082 | Poultry | 2016 | Germany | S. Infantis | PRJEB30317 |
| ERR3263817 | RKI_16-00161 | Human | 2016 | Germany | S. Paratyphi B | PRJEB30317 |
| ERR3263818 | RKI_16-00323 | Human | 2016 | Germany | S. Mbandaka | PRJEB30317 |
| ERR3263819 | RKI_16-00348 | Human | 2016 | Germany | S. Typhi | PRJEB30317 |
| ERR3263820 | RKI_16-00408 | Human | 2016 | Germany | S. Typhi | PRJEB30317 |
| ERR3263821 | RKI_16-00427 | Human | 2016 | Germany | S. Typhi | PRJEB30317 |
| ERR3263822 | RKI_16-00432 | Human | 2016 | Germany | S. Infantis | PRJEB30317 |
| ERR3263823 | RKI_16-00433 | Human | 2016 | Germany | S. Infantis | PRJEB30317 |
| ERR3263824 | RKI_16-00434 | Human | 2016 | Germany | S. Infantis | PRJEB30317 |
| ERR3263825 | RKI_16-00435 | Human | 2016 | Germany | S. Infantis | PRJEB30317 |
| ERR3263826 | RKI_16-00449 | Human | 2016 | Germany | S. Infantis | PRJEB30317 |
| ERR3263827 | RKI_16-00470 | Cattle | 2016 | Germany | S. Infantis | PRJEB30317 |
| ERR3263828 | RKI_16-00471 | Cattle | 2016 | Germany | S. Infantis | PRJEB30317 |
| ERR3263829 | RKI_16-00489 | Human | 2016 | Germany | S. Typhi | PRJEB30317 |
| ERR3263830 | RKI_16-00522 | Human | 2016 | Germany | S. Infantis | PRJEB30317 |
| ERR3263831 | RKI_16-00523 | Pigeon | 2016 | Germany | S. Infantis | PRJEB30317 |
| ERR3263832 | RKI_16-00524 | Human | 2016 | Germany | S. Infantis | PRJEB30317 |
| ERR3263833 | RKI_16-00558 | Human | 2016 | Germany | S. Mbandaka | PRJEB30317 |
| ERR3263834 | RKI_16-00770 | Human | 2016 | Germany | S. Paratyphi B | PRJEB30317 |
| ERR3263835 | RKI_16-00904 | Human | 2016 | Germany | S. Typhi | PRJEB30317 |
| ERR3263836 | RKI_16-00926 | Human | 2016 | Germany | S. Choleraesuis | PRJEB30317 |
| ERR3263837 | RKI_16-00953 | Human | 2016 | Germany | S. Choleraesuis | PRJEB30317 |
| ERR3263838 | RKI_16-00969 | Human | 2016 | Germany | S. Typhi | PRJEB30317 |
| ERR3263839 | RKI_16-01113 | Human | 2016 | Germany | S. Enteritidis | PRJEB30317 |
| ERR3263840 | RKI_16-01218 | Feed | 2016 | Germany | S. Mbandaka | PRJEB30317 |
| ERR3263841 | RKI_16-01366 | Human | 2016 | Germany | S. Enteritidis | PRJEB30317 |
| ERR3263842 | RKI_16-01381 | Human | 2016 | Germany | S. Enteritidis | PRJEB30317 |

| | | | | | | |
|---|---|---|---|---|---|---|
| ERR3263843 | RKI_16-01426 | Human | 2016 | Germany | S. Mbandaka | PRJEB30317 |
| ERR3263844 | RKI_16-01427 | Human | 2016 | Germany | S. Mbandaka | PRJEB30317 |
| ERR3263845 | RKI_16-01428 | Human | 2016 | Germany | S. Mbandaka | PRJEB30317 |
| ERR3263846 | RKI_16-01434 | Human | 2016 | Germany | S. Mbandaka | PRJEB30317 |
| ERR3263847 | RKI_16-01435 | Human | 2016 | Germany | S. Mbandaka | PRJEB30317 |
| ERR3263848 | RKI_16-01473 | Human | 2016 | Germany | S. Mbandaka | PRJEB30317 |
| ERR3263849 | RKI_16-01474 | Human | 2016 | Germany | S. Mbandaka | PRJEB30317 |
| ERR3263850 | RKI_16-01475 | Human | 2016 | Germany | S. Mbandaka | PRJEB30317 |
| ERR3263851 | RKI_16-01476 | Human | 2016 | Germany | S. Mbandaka | PRJEB30317 |
| ERR3263852 | RKI_16-01477 | Human | 2016 | Germany | S. Mbandaka | PRJEB30317 |
| ERR3263853 | RKI_16-01486 | Human | 2016 | Germany | S. Mbandaka | PRJEB30317 |
| ERR3263854 | RKI_16-01525 | Human | 2016 | Germany | S. Enteritidis | PRJEB30317 |
| ERR3263855 | RKI_16-01538 | Human | 2016 | Germany | S. Enteritidis | PRJEB30317 |
| ERR3263856 | RKI_16-01565 | Layer Eggs | 2016 | Germany | S. Enteritidis | PRJEB30317 |
| ERR3263857 | RKI_16-01577 | Human | 2016 | Germany | S. Enteritidis | PRJEB30317 |
| ERR3263858 | RKI_16-01690 | Human | 2016 | Germany | S. Typhi | PRJEB30317 |
| ERR3263859 | RKI_16-01691 | Human | 2016 | Germany | S. Typhi | PRJEB30317 |
| ERR3263860 | RKI_16-01696 | Human | 2016 | Germany | S. Enteritidis | PRJEB30317 |
| ERR3263861 | RKI_16-01721 | Human | 2016 | Germany | S. Typhi | PRJEB30317 |
| ERR3263862 | RKI_16-01747 | Human | 2016 | Germany | S. Typhi | PRJEB30317 |
| ERR3263863 | RKI_16-01811 | Human | 2016 | Germany | S. Choleraesuis | PRJEB30317 |
| ERR3263864 | RKI_16-01836 | Human | 2016 | Germany | S. Typhi | PRJEB30317 |
| ERR3263865 | RKI_16-01876 | Human | 2016 | Germany | S. Enteritidis | PRJEB30317 |
| ERR3263866 | RKI_16-01895 | Human | 2016 | Germany | S. Choleraesuis | PRJEB30317 |
| ERR3263867 | RKI_16-01922 | Human | 2016 | Germany | S. Enteritidis | PRJEB30317 |
| ERR3263868 | RKI_16-01948 | Human | 2016 | Germany | S. Paratyphi B | PRJEB30317 |
| ERR3263869 | RKI_16-01971 | Human | 2016 | Germany | S. Typhimurium | PRJEB30317 |
| ERR3263870 | RKI_16-01981 | Human | 2016 | Germany | S. Enteritidis | PRJEB30317 |
| ERR3263871 | RKI_16-01999 | Human | 2016 | Germany | S. Enteritidis | PRJEB30317 |
| ERR3263872 | RKI_16-02023 | Human | 2016 | Germany | S. Enteritidis | PRJEB30317 |
| ERR3263873 | RKI_16-02033 | Human | 2016 | Germany | S. Enteritidis | PRJEB30317 |
| ERR3263874 | RKI_16-02037 | Human | 2016 | Germany | S. Enteritidis | PRJEB30317 |
| ERR3263875 | RKI_16-02089 | Human | 2016 | Germany | S. ssp.I serol. Rough | PRJEB30317 |
| ERR3263876 | RKI_16-02098 | Human | 2016 | Germany | S. Enteritidis | PRJEB30317 |
| ERR3263877 | RKI_16-02099 | Human | 2016 | Germany | S. Typhi | PRJEB30317 |
| ERR3263878 | RKI_16-02100 | Human | 2016 | Germany | S. Typhi | PRJEB30317 |
| ERR3263879 | RKI_16-02155 | Human | 2016 | Germany | S. Typhi | PRJEB30317 |
| ERR3263880 | RKI_16-02195 | Human | 2016 | Germany | S. ssp.I serol. Rough | PRJEB30317 |
| ERR3263881 | RKI_16-02229 | Human | 2016 | Germany | S. Choleraesuis | PRJEB30317 |
| ERR3263882 | RKI_16-02283 | Human | 2016 | Germany | S. Enteritidis | PRJEB30317 |
| ERR3263883 | RKI_16-02376 | Human | 2016 | Germany | S. Enteritidis | PRJEB30317 |
| ERR3263884 | RKI_16-02387 | Human | 2016 | Germany | S. Enteritidis | PRJEB30317 |
| ERR3263885 | RKI_16-02408 | Human | 2016 | Germany | S. Enteritidis | PRJEB30317 |
| ERR3263886 | RKI_16-02411 | Human | 2016 | Germany | S. Enteritidis | PRJEB30317 |
| ERR3263887 | RKI_16-02549 | Human | 2016 | Germany | S. Typhi | PRJEB30317 |
| ERR3263888 | RKI_16-02633 | Not Provide | 2016 | Germany | S. Typhi | PRJEB30317 |
| ERR3263889 | RKI_16-02654 | Human | 2016 | Germany | S. ssp.I serol. Rough | PRJEB30317 |
| ERR3263890 | RKI_16-02709 | Human | 2016 | Germany | S. Infantis | PRJEB30317 |
| ERR3263891 | RKI_16-02763 | Human | 2016 | Germany | S. Enterititdis | PRJEB30317 |
| ERR3263892 | RKI_16-02787 | Human | 2016 | Germany | S. Typhimurium | PRJEB30317 |
| ERR3263893 | RKI_16-02791 | Poultry | 2016 | Germany | S. ssp.I serol. Rough | PRJEB30317 |
| ERR3263894 | RKI_16-02809 | Human | 2016 | Germany | S. ssp.II serol. Rough | PRJEB30317 |
| ERR3263895 | RKI_16-02844 | Human | 2016 | Germany | S. Typhimurium | PRJEB30317 |
| ERR3263896 | RKI_16-02879 | Human | 2016 | Germany | S. Enteritidis | PRJEB30317 |
| ERR3263897 | RKI_16-02880 | Human | 2016 | Germany | S. Enteritidis | PRJEB30317 |
| ERR3263898 | RKI_16-02881 | Human | 2016 | Germany | S. Enteritidis | PRJEB30317 |
| ERR3263899 | RKI_16-02882 | Human | 2016 | Germany | S. Enteritidis | PRJEB30317 |
| ERR3263900 | RKI_16-02883 | Human | 2016 | Germany | S. Enteritidis | PRJEB30317 |
| ERR3263901 | RKI_16-02907 | Human | 2016 | Germany | S. Paratyphi B | PRJEB30317 |
| ERR3263902 | RKI_16-02909 | Human | 2016 | Germany | S. Enteritidis | PRJEB30317 |
| ERR3263903 | RKI_16-02939 | Human | 2016 | Germany | S. Typhimurium | PRJEB30317 |
| ERR3263904 | RKI_16-02940 | Human | 2016 | Germany | S. Enteritidis | PRJEB30317 |
| ERR3263905 | RKI_16-02952 | Human | 2016 | Germany | S. Enteritidis | PRJEB30317 |
| ERR3263906 | RKI_16-02957 | Human | 2016 | Germany | S. Typhimurium | PRJEB30317 |
| ERR3263907 | RKI_16-02966 | Human | 2016 | Germany | S. Stourbridge | PRJEB30317 |
| ERR3263908 | RKI_16-02983 | Human | 2016 | Germany | S. Enteritidis | PRJEB30317 |
| ERR3263909 | RKI_16-02986 | Human | 2016 | Germany | S. Enteritidis | PRJEB30317 |
| ERR3263910 | RKI_16-02987 | Human | 2016 | Germany | S. Stourbridge | PRJEB30317 |
| ERR3263911 | RKI_16-02988 | Human | 2016 | Germany | S. Stourbridge | PRJEB30317 |
| ERR3263912 | RKI_16-03006 | Human | 2016 | Germany | S. Typhimurium | PRJEB30317 |
| ERR3263913 | RKI_16-03008 | Human | 2016 | Germany | S. Enteritidis | PRJEB30317 |
| ERR3263914 | RKI_16-03038 | Human | 2016 | Germany | S. Enteritidis | PRJEB30317 |
| ERR3263915 | RKI_16-03039 | Human | 2016 | Germany | S. Typhi | PRJEB30317 |
| ERR3263916 | RKI_16-03041 | Human | 2016 | Germany | S. Enteritidis | PRJEB30317 |
| ERR3263917 | RKI_16-03067 | Human | 2016 | Germany | S. Typhimurium | PRJEB30317 |
| ERR3263918 | RKI_16-03092 | Human | 2016 | Germany | S. Typhimurium | PRJEB30317 |

| | | | | | | |
|---|---|---|---|---|---|---|
| ERR3263919 | RKI_16-03094 | Human | 2016 | Germany | S. Enteritidis | PRJEB30317 |
| ERR3263920 | RKI_16-03112 | Human | 2016 | Germany | S. Enteritidis | PRJEB30317 |
| ERR3263921 | RKI_16-03113 | Human | 2016 | Germany | S. Enteritidis | PRJEB30317 |
| ERR3263922 | RKI_16-03117 | Human | 2016 | Germany | S. Enteritidis | PRJEB30317 |
| ERR3263923 | RKI_16-03123 | Human | 2016 | Germany | S. Typhimurium | PRJEB30317 |
| ERR3263924 | RKI_16-03124 | Human | 2016 | Germany | S. Typhimurium | PRJEB30317 |
| ERR3263925 | RKI_16-03151 | Human | 2016 | Germany | S. Enteritidis | PRJEB30317 |
| ERR3263926 | RKI_16-03155 | Human | 2016 | Germany | S. Typhimurium | PRJEB30317 |
| ERR3263927 | RKI_16-03250 | Human | 2016 | Germany | S. Typhimurium | PRJEB30317 |
| ERR3263928 | RKI_16-03252 | Human | 2016 | Germany | S. Enteritidis | PRJEB30317 |
| ERR3263929 | RKI_16-03338 | Human | 2016 | Germany | S. Typhi | PRJEB30317 |
| ERR3263930 | RKI_16-03421 | Human | 2016 | Germany | S. Enteritidis | PRJEB30317 |
| ERR3263931 | RKI_16-03432 | Human | 2016 | Germany | S. Enteritidis | PRJEB30317 |
| ERR3263932 | RKI_16-03433 | Human | 2016 | Germany | S. Enteritidis | PRJEB30317 |
| ERR3263933 | RKI_16-03434 | Human | 2016 | Germany | S. Enteritidis | PRJEB30317 |
| ERR3263934 | RKI_16-03435 | Human | 2016 | Germany | S. Enteritidis | PRJEB30317 |
| ERR3263935 | RKI_16-03436 | Human | 2016 | Germany | S. Enteritidis | PRJEB30317 |
| ERR3263936 | RKI_16-03437 | Human | 2016 | Germany | S. Enteritidis | PRJEB30317 |
| ERR3263937 | RKI_16-03446 | Human | 2016 | Germany | S. Infantis | PRJEB30317 |
| ERR3263938 | RKI_16-03451 | Human | 2016 | Germany | S. Stourbridge | PRJEB30317 |
| ERR3263939 | RKI_16-03453 | Human | 2016 | Germany | S. Enteritidis | PRJEB30317 |
| ERR3263940 | RKI_16-03454 | Human | 2016 | Germany | S. Enteritidis | PRJEB30317 |
| ERR3263941 | RKI_16-03455 | Human | 2016 | Germany | S. Enteritidis | PRJEB30317 |
| ERR3263942 | RKI_16-03456 | Human | 2016 | Germany | S. Enteritidis | PRJEB30317 |
| ERR3263943 | RKI_16-03457 | Human | 2016 | Germany | S. Enteritidis | PRJEB30317 |
| ERR3263944 | RKI_16-03458 | Human | 2016 | Germany | S. Enteritidis | PRJEB30317 |
| ERR3263945 | RKI_16-03459 | Human | 2016 | Germany | S. Enteritidis | PRJEB30317 |
| ERR3263946 | RKI_16-03493 | Human | 2016 | Germany | S. Infantis | PRJEB30317 |
| ERR3263947 | RKI_16-03528 | Human | 2016 | Germany | S. Infantis | PRJEB30317 |
| ERR3263948 | RKI_16-03529 | Human | 2016 | Germany | S. Typhimurium | PRJEB30317 |
| ERR3263949 | RKI_16-03540 | Human | 2016 | Germany | S. Typhi | PRJEB30317 |
| ERR3263950 | RKI_16-03559 | Human | 2016 | Germany | S. Infantis | PRJEB30317 |
| ERR3263951 | RKI_16-03637 | Human | 2016 | Germany | S. Typhi | PRJEB30317 |
| ERR3263952 | RKI_16-03721 | Human | 2016 | Germany | S. Typhi | PRJEB30317 |
| ERR3263953 | RKI_16-03723 | Human | 2016 | Germany | S. Kentucky | PRJEB30317 |
| ERR3263954 | RKI_16-03739 | Human | 2016 | Germany | S. Typhimurium | PRJEB30317 |
| ERR3263955 | RKI_16-03748 | Human | 2016 | Germany | S. Infantis | PRJEB30317 |
| ERR3263956 | RKI_16-03767 | Human | 2016 | Germany | S. Infantis | PRJEB30317 |
| ERR3263957 | RKI_16-03838 | Human | 2016 | Germany | S. Infantis | PRJEB30317 |
| ERR3263958 | RKI_16-03841 | Human | 2016 | Germany | S. Infantis | PRJEB30317 |
| ERR3263959 | RKI_16-03855 | Human | 2016 | Germany | S. Infantis | PRJEB30317 |
| ERR3263960 | RKI_16-03900 | Human | 2016 | Germany | S. Infantis | PRJEB30317 |
| ERR3263961 | RKI_16-03904 | Human | 2016 | Germany | S. Stourbridge | PRJEB30317 |
| ERR3263962 | RKI_16-03913 | Human | 2016 | Germany | S. Typhi | PRJEB30317 |
| ERR3263963 | RKI_16-03921 | Human | 2016 | Germany | S. Stourbridge | PRJEB30317 |
| ERR3263964 | RKI_16-03940 | Human | 2016 | Germany | S. Typhi | PRJEB30317 |
| ERR3263965 | RKI_16-03949 | Human | 2016 | Germany | S. Infantis | PRJEB30317 |
| ERR3263966 | RKI_16-03953 | Human | 2016 | Germany | S. Infantis | PRJEB30317 |
| ERR3263967 | RKI_16-03954 | Human | 2016 | Germany | S. Infantis | PRJEB30317 |
| ERR3263968 | RKI_16-03970 | Human | 2016 | Germany | S. Infantis | PRJEB30317 |
| ERR3263969 | RKI_16-03971 | Human | 2016 | Germany | S. Infantis | PRJEB30317 |
| ERR3263970 | RKI_16-03972 | Human | 2016 | Germany | S. Infantis | PRJEB30317 |
| ERR3263971 | RKI_16-03980 | Human | 2016 | Germany | S. Stourbridge | PRJEB30317 |
| ERR3263972 | RKI_16-04059 | Human | 2016 | Germany | S. Typhimurium | PRJEB30317 |
| ERR3263973 | RKI_16-04083 | Human | 2016 | Germany | S. Typhimurium | PRJEB30317 |
| ERR3263974 | RKI_16-04084 | Human | 2016 | Germany | S. Infantis | PRJEB30317 |
| ERR3263975 | RKI_16-04173 | Human | 2016 | Germany | S. Typhi | PRJEB30317 |
| ERR3263976 | RKI_16-04195 | Human | 2016 | Germany | S. Typhi | PRJEB30317 |
| ERR3263977 | RKI_16-04218 | Human | 2016 | Germany | S. Typhimurium | PRJEB30317 |
| ERR3263978 | RKI_16-04298 | Human | 2016 | Germany | S. Stourbridge | PRJEB30317 |
| ERR3263979 | RKI_16-04300 | Human | 2016 | Germany | S. Typhimurium | PRJEB30317 |
| ERR3263980 | RKI_16-04315 | Human | 2016 | Germany | S. Kentucky | PRJEB30317 |
| ERR3263981 | RKI_16-04352 | Human | 2016 | Germany | S. Choleraesuis | PRJEB30317 |
| ERR3263982 | RKI_16-04353 | Human | 2016 | Germany | S. Choleraesuis | PRJEB30317 |
| ERR3263983 | RKI_16-04355 | Human | 2016 | Germany | S. Choleraesuis | PRJEB30317 |
| ERR3263984 | RKI_16-04356 | Human | 2016 | Germany | S. Choleraesuis | PRJEB30317 |
| ERR3263985 | RKI_16-04357 | Human | 2016 | Germany | S. Choleraesuis | PRJEB30317 |
| ERR3263986 | RKI_16-04375 | Human | 2016 | Germany | S. enterica 11:z41:e,n,z15 | PRJEB30317 |
| ERR3263987 | RKI_16-04383-1 | Human | 2016 | Germany | S. Typhi | PRJEB30317 |
| ERR3263988 | RKI_16-04441 | Human | 2016 | Germany | S. Typhi | PRJEB30317 |
| ERR3263989 | RKI_16-04466 | Human | 2016 | Germany | S. Choleraesuis | PRJEB30317 |
| ERR3263990 | RKI_16-04486 | Human | 2016 | Germany | S. Infantis | PRJEB30317 |
| ERR3263991 | RKI_16-04495 | Human | 2016 | Germany | S. Infantis | PRJEB30317 |
| ERR3263992 | RKI_16-04501 | Human | 2016 | Germany | S. Infantis | PRJEB30317 |
| ERR3263993 | RKI_16-04605 | Human | 2016 | Germany | S. Typhi | PRJEB30317 |
| ERR3263994 | RKI_16-04615 | Human | 2016 | Germany | S. Typhi | PRJEB30317 |

| | | | | | | |
|---|---|---|---|---|---|---|
| ERR3263995 | RKI_16-04676 | Human | 2016 | Germany | S. Typhi | PRJEB30317 |
| ERR3263996 | RKI_16-04685 | Human | 2016 | Germany | S. Choleraesuis | PRJEB30317 |
| ERR3263997 | RKI_16-04832 | Human | 2016 | Germany | S. Choleraesuis | PRJEB30317 |
| ERR3263998 | RKI_16-04870 | Human | 2016 | Germany | S. Choleraesuis | PRJEB30317 |
| ERR3263999 | RKI_16-04871 | Human | 2016 | Germany | S. Choleraesuis | PRJEB30317 |
| ERR3264000 | RKI_16-04876 | Human | 2016 | Germany | S. Typhi | PRJEB30317 |
| ERR3264001 | RKI_16-04907 | Human | 2016 | Germany | S. Choleraesuis var. Kunzendorf (mono) | PRJEB30317 |
| ERR3264002 | RKI_16-04910 | Human | 2016 | Germany | S. Enteritidis | PRJEB30317 |
| ERR3264003 | RKI_16-04913 | Human | 2016 | Germany | S. Typhimurium | PRJEB30317 |
| ERR3264004 | RKI_16-04930 | Human | 2016 | Germany | S. Choleraesuis | PRJEB30317 |
| ERR3264005 | RKI_16-04936 | Human | 2016 | Germany | S. Typhimurium | PRJEB30317 |
| ERR3264006 | RKI_16-04938 | Human | 2016 | Germany | S. Typhimurium | PRJEB30317 |
| ERR3264007 | RKI_16-04939 | Human | 2016 | Germany | S. Typhimurium | PRJEB30317 |
| ERR3264008 | RKI_16-04940 | Human | 2016 | Germany | S. Typhimurium | PRJEB30317 |
| ERR3264009 | RKI_16-04961 | Human | 2016 | Germany | S. Choleraesuis | PRJEB30317 |
| ERR3264010 | RKI_18-06929 | Human | 2018 | Germany | S. Typhi | PRJEB30317 |
| ERR3264011 | RKI_16-05058 | Human | 2016 | Germany | S. enterica 11:z41:e,n,z15 | PRJEB30317 |
| ERR3264012 | RKI_16-05098 | Human | 2016 | Germany | S. Typhi | PRJEB30317 |
| ERR3264013 | RKI_16-05374 | Human | 2016 | Germany | S. Choleraesuis | PRJEB30317 |
| ERR3264014 | RKI_16-05392 | Human | 2016 | Germany | S. Choleraesuis | PRJEB30317 |
| ERR3264015 | RKI_16-05399 | Human | 2016 | Germany | S. Choleraesuis | PRJEB30317 |
| ERR3264016 | RKI_17-00006 | Human | 2017 | Germany | S. enterica 11:z41:e,n,z15 | PRJEB30317 |
| ERR3264017 | RKI_17-00007 | Human | 2017 | Germany | S. enterica 11:z41:e,n,z15 | PRJEB30317 |
| ERR3264018 | RKI_17-00008 | Human | 2017 | Germany | S. enterica 11:z41:e,n,z15 | PRJEB30317 |
| ERR3264019 | RKI_17-00036 | Human | 2017 | Germany | S. Choleraesuis | PRJEB30317 |
| ERR3264020 | RKI_17-00045 | Human | 2017 | Germany | S. Choleraesuis | PRJEB30317 |
| ERR3264022 | RKI_17-00254 | Human | 2017 | Germany | S. Paratyphi C | PRJEB30317 |
| ERR3264023 | RKI_17-00280 | Human | 2017 | Germany | S. Choleraesuis | PRJEB30317 |
| ERR3264024 | RKI_17-00296 | Human | 2017 | Germany | S. enterica 11:z41:e,n,z15 | PRJEB30317 |
| ERR3264025 | RKI_17-00390 | Human | 2017 | Germany | S. Choleraesuis | PRJEB30317 |
| ERR3264026 | RKI_17-00391 | Human | 2017 | Germany | S. Choleraesuis var. Kunzendorf (mono) | PRJEB30317 |
| ERR3264027 | RKI_17-00399 | Human | 2017 | Germany | S. Stourbridge | PRJEB30317 |
| ERR3264028 | RKI_17-00449 | Human | 2017 | Germany | S. Stourbridge | PRJEB30317 |
| ERR3264029 | RKI_17-00460 | Human | 2017 | Germany | S. enterica 11:z41:e,n,z15 | PRJEB30317 |
| ERR3264030 | RKI_17-00643 | Human | 2017 | Germany | S. Choleraesuis | PRJEB30317 |
| ERR3264031 | RKI_17-00666 | Human | 2017 | Germany | S. enterica 11:z41:e,n,z15 | PRJEB30317 |
| ERR3264032 | RKI_17-00682 | Human | 2017 | Germany | S. Stourbridge | PRJEB30317 |
| ERR3264033 | RKI_17-00696 | Human | 2017 | Germany | S. Choleraesuis | PRJEB30317 |
| ERR3264034 | RKI_17-00717 | Human | 2017 | Germany | S. Paratyphi C | PRJEB30317 |
| ERR3264035 | RKI_17-00851 | Human | 2017 | Germany | S. Choleraesuis var. Kunzendorf (mono) | PRJEB30317 |
| ERR3264036 | RKI_17-00913 | Human | 2017 | Germany | S. ssp.I serol. Rough | PRJEB30317 |
| ERR3264037 | RKI_17-00942 | Human | 2017 | Germany | S. enterica 11:z41:e,n,z15 | PRJEB30317 |
| ERR3264038 | RKI_17-00957 | Human | 2017 | Germany | S. Choleraesuis | PRJEB30317 |
| ERR3264039 | RKI_17-00963 | Human | 2017 | Germany | S. Stourbridge | PRJEB30317 |
| ERR3264040 | RKI_17-01071 | Human | 2017 | Germany | S. enterica 11:z41:e,n,z15 | PRJEB30317 |
| ERR3264041 | RKI_17-02304 | Human | 2017 | Germany | S. Kentucky | PRJEB30317 |
| ERR3264042 | RKI_17-02411 | Human | 2017 | Germany | S. Kentucky | PRJEB30317 |
| ERR3264043 | RKI_17-02757 | Human | 2017 | Germany | S. Kentucky | PRJEB30317 |
| ERR3264044 | RKI_17-04278 | Human | 2017 | Germany | S. Kottbus | PRJEB30317 |
| ERR3264045 | RKI_17-04279 | Human | 2017 | Germany | S. Kottbus | PRJEB30317 |
| ERR3264046 | RKI_17-04280 | Human | 2017 | Germany | S. Kottbus | PRJEB30317 |
| ERR3264047 | RKI_17-04281 | Human | 2017 | Germany | S. Kottbus | PRJEB30317 |
| ERR3264048 | RKI_17-04282 | Human | 2017 | Germany | S. Kottbus | PRJEB30317 |
| ERR3264049 | RKI_17-04283 | Human | 2017 | Germany | S. Kottbus | PRJEB30317 |
| ERR3264050 | RKI_17-04284 | Human | 2017 | Germany | S. Kottbus | PRJEB30317 |
| ERR3264051 | RKI_17-04285 | Human | 2017 | Germany | S. Kottbus | PRJEB30317 |
| ERR3264052 | RKI_17-04437 | Human | 2017 | Germany | S. Kottbus | PRJEB30317 |
| ERR3264053 | RKI_17-04439 | Human | 2017 | Germany | S. Kottbus | PRJEB30317 |
| ERR3264054 | RKI_17-04795 | Human | 2017 | Germany | S. Kottbus | PRJEB30317 |
| ERR3264055 | RKI_17-04797 | Human | 2017 | Germany | S. Kentucky | PRJEB30317 |
| ERR3264056 | RKI_17-05582 | Human | 2017 | Germany | S. Kottbus | PRJEB30317 |
| ERR3264057 | RKI_17-06287 | Human | 2017 | Germany | S. Agona | PRJEB30317 |
| ERR3264058 | RKI_17-06303 | Human | 2017 | Germany | S. Choleraesuis | PRJEB30317 |
| ERR3264059 | RKI_17-06684 | Human | 2017 | Germany | S. Agona | PRJEB30317 |
| ERR3264060 | RKI_17-06869 | Human | 2017 | Germany | S. Kentucky | PRJEB30317 |
| ERR3264062 | RKI_18-00068 | Human | 2018 | Germany | S. Enteritidis | PRJEB30317 |
| ERR3264063 | RKI_18-00072 | Human | 2018 | Germany | S. Choleraesuis | PRJEB30317 |
| ERR3264064 | RKI_18-00177 | Human | 2018 | Germany | S. Choleraesuis | PRJEB30317 |
| ERR3264065 | RKI_18-00253 | Human | 2018 | Germany | S. Choleraesuis | PRJEB30317 |
| ERR3264066 | RKI_18-00651 | Human | 2018 | Germany | S. Choleraesuis | PRJEB30317 |
| ERR3264067 | RKI_18-00670 | Human | 2018 | Germany | S. Choleraesuis | PRJEB30317 |
| ERR3264068 | RKI_18-00952 | Human | 2018 | Germany | S. Agona | PRJEB30317 |
| ERR3264069 | RKI_18-01921 | Human | 2018 | Germany | S. Derby | PRJEB30317 |
| ERR3264070 | RKI_18-02045 | Human | 2018 | Germany | S. Derby | PRJEB30317 |
| ERR3264071 | RKI_18-02153 | Human | 2018 | Germany | S. Derby | PRJEB30317 |
| ERR3264072 | RKI_18-02673 | Human | 2018 | Germany | S. Derby | PRJEB30317 |

| | | | | | | |
|---|---|---|---|---|---|---|
| ERR3264073 | RKI_18-03486 | Human | 2018 | Germany | S. Derby | PRJEB30317 |
| ERR3264074 | RKI_18-04245 | Human | 2017 | Germany | S. Enteritidis | PRJEB30317 |
| ERR3264075 | RKI_18-04526 | Human | 2018 | Germany | S. Infantis | PRJEB30317 |
| ERR3264076 | RKI_18-04597 | Human | 2018 | Germany | S. Infantis | PRJEB30317 |
| ERR3264077 | RKI_18-04831 | Human | 2018 | Germany | S. Infantis | PRJEB30317 |
| ERR3264078 | RKI_18-04962 | Human | 2018 | Germany | S. Infantis | PRJEB30317 |
| ERR3264079 | RKI_18-05170 | Human | 2018 | Germany | S. Infantis | PRJEB30317 |
| ERR3264080 | RKI_18-05807 | Human | 2018 | Germany | S. Mikawasima | PRJEB30317 |
| ERR3264081 | RKI_18-06149 | Human | 2018 | Germany | S. Mikawasima | PRJEB30317 |
| ERR3264082 | RKI_18-06220 | Human | 2018 | Germany | S. Mikawasima | PRJEB30317 |
| ERR3264083 | RKI_18-06230 | Human | 2018 | Germany | S. Mikawasima | PRJEB30317 |
| ERR3264084 | RKI_18-06286 | Human | 2018 | Germany | S. Mikawasima | PRJEB30317 |
| ERR3264085 | RKI_18-06287 | Human | 2018 | Germany | S. Mikawasima | PRJEB30317 |
| ERR3264086 | RKI_18-06288 | Human | 2018 | Germany | S. Mikawasima | PRJEB30317 |
| ERR3264087 | RKI_18-06449 | Human | 2018 | Germany | S. Mikawasima | PRJEB30317 |
| ERR3264088 | RKI_18-06450 | Human | 2018 | Germany | S. Mikawasima | PRJEB30317 |
| ERR3264089 | RKI_18-06499 | Human | 2018 | Germany | S. Typhimurium | PRJEB30317 |
| ERR3264090 | RKI_18-06654 | Human | 2018 | Germany | S. Mikawasima | PRJEB30317 |
| ERR3264091 | RKI_18-07063 | Human | 2018 | Germany | S. Infantis | PRJEB30317 |
| ERR3264092 | RKI_18-07072 | Human | 2018 | Germany | S. Infantis | PRJEB30317 |
| ERR3264021 | RKI_18-06747 | Human | 2018 | Germany | S. Strathcona | PRJEB30317 |
| ERR3264061 | RKI_18-06757 | Human | 2018 | Germany | S. Strathcona | PRJEB30317 |
| ERR2003327 | RKI_00-10487 | Human | 2000 | Germany | S. Typhimurium mono | PRJEB16326 |
| ERR2003328 | RKI_01-00356 | Human | 2001 | Germany | S. Typhimurium mono | PRJEB16326 |
| ERR2003329 | RKI_01-02645 | Human | 2001 | Germany | S. Typhimurium mono | PRJEB16326 |
| ERR2003330 | RKI_02-02704 | Pigs | 2002 | Germany | S. Typhimurium mono | PRJEB16326 |
| ERR2003331 | RKI_02-06979 | Human | 2002 | Germany | S. Typhimurium mono | PRJEB16326 |
| ERR2003332 | RKI_03-03803 | Pigs | 2003 | Germany | S. Typhimurium mono | PRJEB16326 |
| ERR2003341 | RKI_04-00972 | Human | 2004 | Germany | S. Typhimurium | PRJEB16326 |
| ERR2003333 | RKI_04-00977 | Pigs | 2004 | Germany | S. Typhimurium mono | PRJEB16326 |
| ERR2003334 | RKI_04-05579 | Human | 2004 | Germany | S. Typhimurium mono | PRJEB16326 |
| ERR2003335 | RKI_05-04392 | Cattle | 2005 | Germany | S. Typhimurium mono | PRJEB16326 |
| ERR2099818 | RKI_16-00003 | Human | 2016 | Germany | S. Typhimurium | PRJEB16326 |
| ERR2099823 | RKI_16-00006 | Human | 2016 | Germany | S. Typhimurium | PRJEB16326 |
| ERR2099820 | RKI_16-00021 | Human | 2016 | Germany | S. Typhimurium | PRJEB16326 |
| ERR2099822 | RKI_16-00091 | Pigs | 2016 | Germany | S. Typhimurium | PRJEB16326 |
| ERR1912198 | RKI_16-00803 | Human | 2016 | Germany | S. Typhimurium | PRJEB16326 |
| ERR2003365 | RKI_16-04322 | Human | 2016 | Germany | S. Typhimurium | PRJEB16326 |
| ERR2003364 | RKI_16-04515 | Human | 2016 | Germany | S. Typhimurium mono | PRJEB16326 |
| ERR2003363 | RKI_16-04623 | Human | 2016 | Germany | S. Typhimurium | PRJEB16326 |
| ERR2003362 | RKI_16-04669 | Pigs | 2016 | Germany | S. Typhimurium mono | PRJEB16326 |
| ERR2003361 | RKI_16-04673 | Human | 2016 | Germany | S. Typhimurium mono | PRJEB16326 |
| ERR2003360 | RKI_16-04674 | Human | 2016 | Germany | S. Typhimurium mono | PRJEB16326 |
| ERR2003359 | RKI_16-04689 | Human | 2016 | Germany | S. Typhimurium | PRJEB16326 |
| ERR2003358 | RKI_16-04690 | Human | 2016 | Germany | S. Typhimurium | PRJEB16326 |
| ERR2003357 | RKI_16-04699 | Human | 2016 | Germany | S. Typhimurium | PRJEB16326 |
| ERR2003356 | RKI_16-04700 | Human | 2016 | Germany | S. Typhimurium | PRJEB16326 |
| ERR2003355 | RKI_16-04701 | Human | 2016 | Germany | S. Typhimurium | PRJEB16326 |
| ERR2003354 | RKI_16-04702 | Human | 2016 | Germany | S. Typhimurium | PRJEB16326 |
| ERR2003353 | RKI_16-04715 | Human | 2016 | Germany | S. Typhimurium mono | PRJEB16326 |
| ERR2003352 | RKI_16-04774 | Human | 2016 | Germany | S. Typhimurium | PRJEB16326 |
| ERR2003351 | RKI_16-04783 | Human | 2016 | Germany | S. Typhimurium mono | PRJEB16326 |
| ERR2003350 | RKI_16-04785 | Human | 2016 | Germany | S. Typhimurium mono | PRJEB16326 |
| ERR2003349 | RKI_16-04787 | Human | 2016 | Germany | S. Typhimurium mono | PRJEB16326 |
| ERR2003348 | RKI_16-04803 | Human | 2016 | Germany | S. Typhimurium | PRJEB16326 |
| ERR2003347 | RKI_16-04810 | Human | 2016 | Germany | S. Typhimurium mono | PRJEB16326 |
| ERR2003346 | RKI_16-04828 | Human | 2016 | Germany | S. Typhimurium | PRJEB16326 |
| ERR2003345 | RKI_16-04829 | Human | 2016 | Germany | S. Typhimurium mono | PRJEB16326 |
| ERR2003344 | RKI_16-04854 | Human | 2016 | Germany | S. Typhimurium | PRJEB16326 |
| ERR2003343 | RKI_16-04857 | Human | 2016 | Germany | S. Typhimurium | PRJEB16326 |
| ERR2003342 | RKI_16-04869 | Human | 2016 | Germany | S. Typhimurium mono | PRJEB16326 |
| ERR1950104 | RKI_16-04996 | Human | 2016 | Germany | S. Typhimurium mono | PRJEB16326 |
| ERR2003340 | RKI_97-02972 | Human | 1997 | Germany | S. Typhimurium | PRJEB16326 |
| ERR2003336 | RKI_97-04254 | Human | 1997 | Germany | S. Typhimurium | PRJEB16326 |
| ERR2003337 | RKI_97-04255 | Human | 1997 | Germany | S. Typhimurium | PRJEB16326 |
| ERR2003338 | RKI_97-05971 | Human | 1997 | Germany | S. Typhimurium | PRJEB16326 |
| ERR2003339 | RKI_97-05974 | Human | 1997 | Germany | S. Typhimurium | PRJEB16326 |
